# Cell cycle corruption in a pre-leukemic ETV6-RUNX1 model exposes RUNX1 addiction as a therapeutic target in acute lymphoblastic leukemia

**DOI:** 10.1101/2020.12.22.423823

**Authors:** Jason P Wray, Elitza M Deltcheva, Charlotta Boiers, Simon E Richardson, Jyoti Bikram Chhetri, Sladjana Gagrica, Yanping Guo, Anuradha Illendula, Joost HA Martens, Hendrik G Stunnenberg, John H Bushweller, Rachael Nimmo, Tariq Enver

## Abstract

The ETV6-RUNX1 onco-fusion arises *in utero*, initiating a clinically silent pre-leukemic state associated with the development of pediatric B-acute lymphoblastic leukemia (B-ALL). We characterize the ETV6-RUNX1 regulome by integrating chromatin immunoprecipitation- and RNA-sequencing and show that ETV6-RUNX1 functions primarily through competition for RUNX1 binding sites and transcriptional repression. In pre-leukemia, this results in ETV6-RUNX1 antagonization of cell cycle regulation by RUNX1 as evidenced by mass cytometry analysis of B-lineage cells derived from ETV6-RUNX1 knock-in human pluripotent stem cells. In frank leukemia, knockdown of RUNX1 or its co-factor CBFβ results in cell death suggesting sustained requirement for RUNX1 activity which is recapitulated by chemical perturbation using an allosteric CBFβ-inhibitor. Strikingly, we show that RUNX1 addiction extends to other genetic subtypes of pediatric B-ALL and also adult disease. Importantly, inhibition of RUNX1 activity spares normal hematopoiesis. Our results implicate chemical intervention in the RUNX1 program as an exciting therapeutic opportunity in ALL.

## Introduction

Childhood B-acute lymphoblastic leukaemia (B-ALL) is the most common pediatric cancer and is clinically distinct from adult ALL, characterized by a distinct mutational spectrum, higher incidence and overall good prognosis. However, despite high cure rates in low risk cases, B-ALL remains a leading cause of cancer-related death in children. Current chemotherapy regimens are highly toxic, associated with severe long-term sequelae and relapse occurs in ∼20% of patients. While immunotherapeutic approaches have contributed to improved outcomes, toxicity and relapse remain significant concerns.

Most pediatric leukemias are likely to originate during embryonic and fetal development and are often associated with chromosomal rearrangements that disrupt the normal function of hematopoietic transcription factors (Greaves, 2018). In B-ALL, a paradigmatic example of an oncogene arising *in utero* is the ETS translocation variant 6 (ETV6) - runt-related transcription factor 1 (RUNX1) fusion, affecting two major hematopoietic regulators. ETV6-RUNX1 accounts for ∼25% of pediatric B-ALL cases and is detected in 1-5% of all newborns, as demonstrated by studies of neonatal blood spots and PCR of ligated breakpoints (Greaves and Wiemels, 2003, Schafer et al., 2018). However, the translocation is not sufficient to induce clinically overt disease and only a small proportion of children carrying the fusion will transition from the clinically silent pre-leukaemic state into full blown leukemia. This transition is dependent on the acquisition of secondary mutations that evolve in a branching evolutionary pattern and contribute to the genetic heterogeneity of B-ALL, a confounding factor for both conventional and targeted therapeutic approaches. While key second-hit mutations in ETV6-RUNX1^+^ leukemia involve loss of *CDKN2A* and the second copy of *ETV6*, the fusion gene and the native RUNX1 allele are rarely lost, suggesting that the ETV6-RUNX1 - RUNX1 axis may be required for leukemic maintenance (Sulong et al., 2009, Tsuzuki et al., 2007, Mullighan et al., 2007).

Structurally, ETV6-RUNX1 fuses almost the entire RUNX1 protein to the N-terminal portion of ETV6 and has been proposed to recruit repressors to RUNX1 target genes (Hiebert et al., 1996, Guidez et al., 2000, Hiebert et al., 2001, Morrow et al., 2007, Wiemels and Greaves, 1999). Several studies have examined the function of ETV6-RUNX1 by characterizing its targets in murine or human cell lines (Linka et al., 2013, Teppo et al., 2016, Fuka et al., 2011). Whilst providing useful insights, these do not address the role of ETV6-RUNX1 as a “first-hit” in a pre-leukemic context and in the absence of second hit mutations. Furthermore, while multiple detailed studies have interrogated the relationship between native RUNX1 and the RUNX1-ETO fusion protein in acute myeloid leukemia (AML), such analyses are lacking in B-ALL (Ben-Ami et al., 2013, Ptasinska et al., 2014).

RUNX1 and its co-factor CBFβ, together comprising the <u>C</u>ore <u>B</u>inding <u>F</u>actor (CBF) complex, are the most frequent targets of mutations in hematological malignancies. Genetic changes affecting RUNX1, such as translocations or point mutations, are generally linked to loss of function, broadly classifying it as a tumor suppressor (Osato, 2004). Notably, these mutations are usually heterozygous and the second, wild-type copy is rarely lost and often amplified, suggesting that retention of RUNX1 might offer a selective advantage for leukemic cells (Mikhail et al., 2002, Mullighan, 2012, Attarbaschi et al., 2004). Recent reports have indeed shown a supporting role for RUNX1 in leukemogenesis, primarily in AML, but also in T-ALL and mixed-lineage leukemia (MLL), offering a therapeutic rationale for targeting RUNX1 and its cofactor CBFβ (Wilkinson et al., 2013, Choi et al., 2017, Ben-Ami et al., 2013, Ptasinska et al., 2014). Accordingly, several structural derivatives of an allosteric CBFβ inhibitor have been tested in these diseases, with promising results (Illendula et al., 2016, Choi et al., 2017). It remains to be assessed if B-ALL cells are dependent on RUNX1, but its requirement for the survival of early mouse B-cell progenitors suggests that its inhibition may represent a therapeutic opportunity (Niebuhr et al., 2013).

Here, we characterize the ETV6-RUNX1 chimeric- and native RUNX1-responsive regulomes and explore their relationship in the context of pre-leukemia and overt disease. Using mass cytometry we functionally demonstrate that as a “first-hit” ETV6-RUNX1 alters cell cycle of early B-cell progenitors by hijacking and repressing RUNX1 targets. The functional antagonism between the two transcription factors exposes an addiction to RUNX1-activity in overt leukaemia, spanning pediatric and adult B-ALL subtypes. We demonstrate that this vulnerability has therapeutic potential and can be exploited using an allosteric inhibitor of the CBF complex.

## Results

### Delineating the ETV6-RUNX1 regulome in childhood B-ALL

To identify direct targets of ETV6-RUNX1 we performed ChIP-seq in the cell line Reh, a widely used model system for t(12;21) ALL. As Reh lacks the second ETV6 allele, ETV6-RUNX1 can be specifically immunoprecipitated using ETV6 antibodies. Considering only peaks identified with two independent ETV6 antibodies and overlapping DNaseI-hypersensitivity sites, 1171 high confidence binding sites were identified (Figure 1A, upper panel, marked with red dashed line, Figure S1A). These peaks were highly enriched for motifs including RUNX, ETS, Ascl2 (NHLH1) and SP1 (Figure 1B, Table S1).

**Figure 1.**
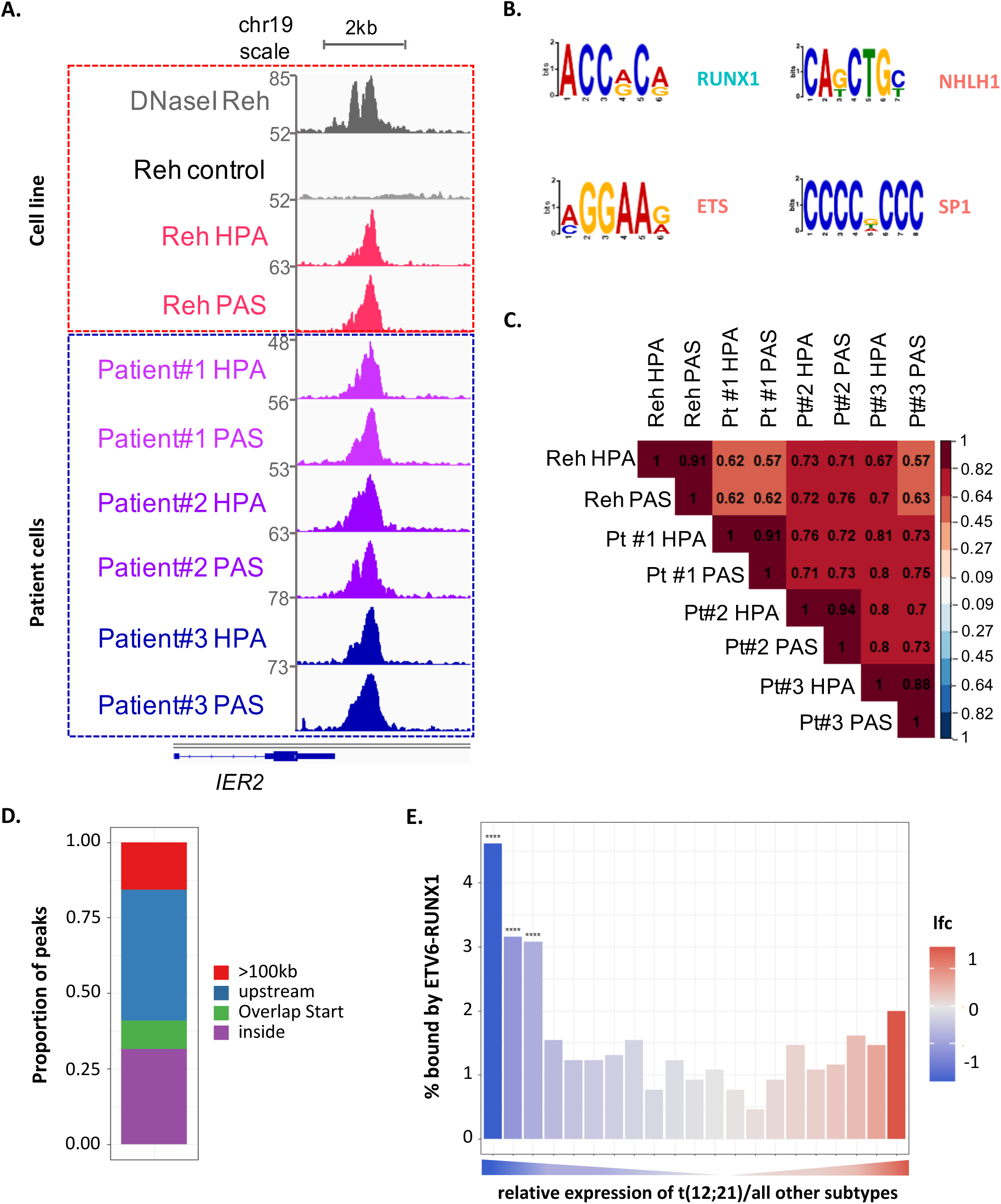
Delineating the ETV6-RUNX1 regulome in childhood B-ALL. (A) Integrative Genomics Viewer (IGV) screenshot of ETV6-RUNX1 binding at the IER2 locus (Lachmann et al., 2010) in an ETV6-RUNX1^+^ cell line (Reh, upper panel, red dashed line) or patient-derived ETV6-RUNX1^+^ leukaemia samples (bottom panel, blue dashed line). HPA and PAS, two independent ETV6 antibodies; DNaseI, DNaseI hypersensitivity sequencing; control, signal following immunoprecipitation with an IgG antibody. (B) MEME enrichment of motifs in sites bound by ETV6-RUNX1. See also Table S1 (enrichment motifs). (C) Correlation matrix for normalized signal across all ETV6-RUNX1 binding sites. (D) Distribution of peaks relative to their nearest genes (see methods for analysis). (E) ETV6-RUNX1 binding across genes binned according to relative expression in ETV6-RUNX1^+^ B-ALL as compared to all other subtypes. **** p<0.0001, hypergeometric test.

To validate the clinical relevance of the peaks, we performed ETV6 ChIP-seq in three t(12;21) patient samples with low or undetectable ETV6 expression and compared their binding patterns to Reh cells (Figure 1A, bottom panel, marked with blue dashed line, Figure S1B). Notably, our analysis revealed a strong correlation across datasets with the majority of high-confidence sites identified in Reh cells also found in patients (Figure 1C). Peaks detected in all eight samples (711 in total) had the highest binding affinity (Figure S1C) with the majority found to be intragenic or upstream of the TSS (Figure 1D). Motif analysis confirmed a preference for canonical RUNX and ETS motifs with a prevalence of RUNX-motif in the higher affinity peaks, consistent with ETV6-RUNX1 interacting with chromatin primarily through the runt domain (Figure S1D and S1E).

We next examined the relationship between ETV6-RUNX1 target binding and transcriptional regulation, taking advantage of publicly available transcriptome datasets from diagnostic childhood B-ALLs (Gu et al., 2019). We compared ETV6-RUNX1^+^ samples to all other subtypes, ranking genes according to their relative expression. A trend towards increased ETV6-RUNX1 binding was observed regardless of expression directionality with a statistically significant enrichment for downregulated genes (Figure 1E).

These data suggest that in the patient setting ETV6-RUNX1 activity is primarily associated with repression.

### *ETV6-RUNX1* induces transcriptional changes indicative of cell cycle repression in pre-leukemia

In order to link ETV6-RUNX1 genome occupancy with transcriptional regulation, we next performed RNA-seq following knockdown of the fusion gene in Reh cells using two independent shRNAs (Figure S2A). Differentially expressed genes (DEGs) were found to be significantly enriched for ETV6-RUNX1 binding, with a prevalence of upregulated genes (Figure 2A, left panel). As ETV6-RUNX1 is an initiating event arising *in utero*, its initial impact is on a fetal cell in the absence of additional mutations acquired during leukemic progression. To explore this, we made use of RNA-seq data from the hIPSC ETV6-RUNX1 knock-in system developed in our lab, which displays a partial block in B-cell differentiation (Figures S2B and S2C) (Boiers et al., 2018). iPS derived ETV6-RUNX1^+^ proB cells showed significant enrichment for ETV6-RUNX1 binding among the DEGs, with a prevalence of downregulated genes (Figures 2A, right panel). In addition, DEGs from the two systems were strongly anticorrelated demonstrating the preservation of the ETV6-RUNX1 regulome across these model systems (Figure 2B). Together, these data show that in both overt and pre-leukemic contexts ETV6-RUNX1 acts predominantly, but not exclusively, as a repressor in agreement with our analysis of the patient cohort (Figure 1E).

**Figure 2.**
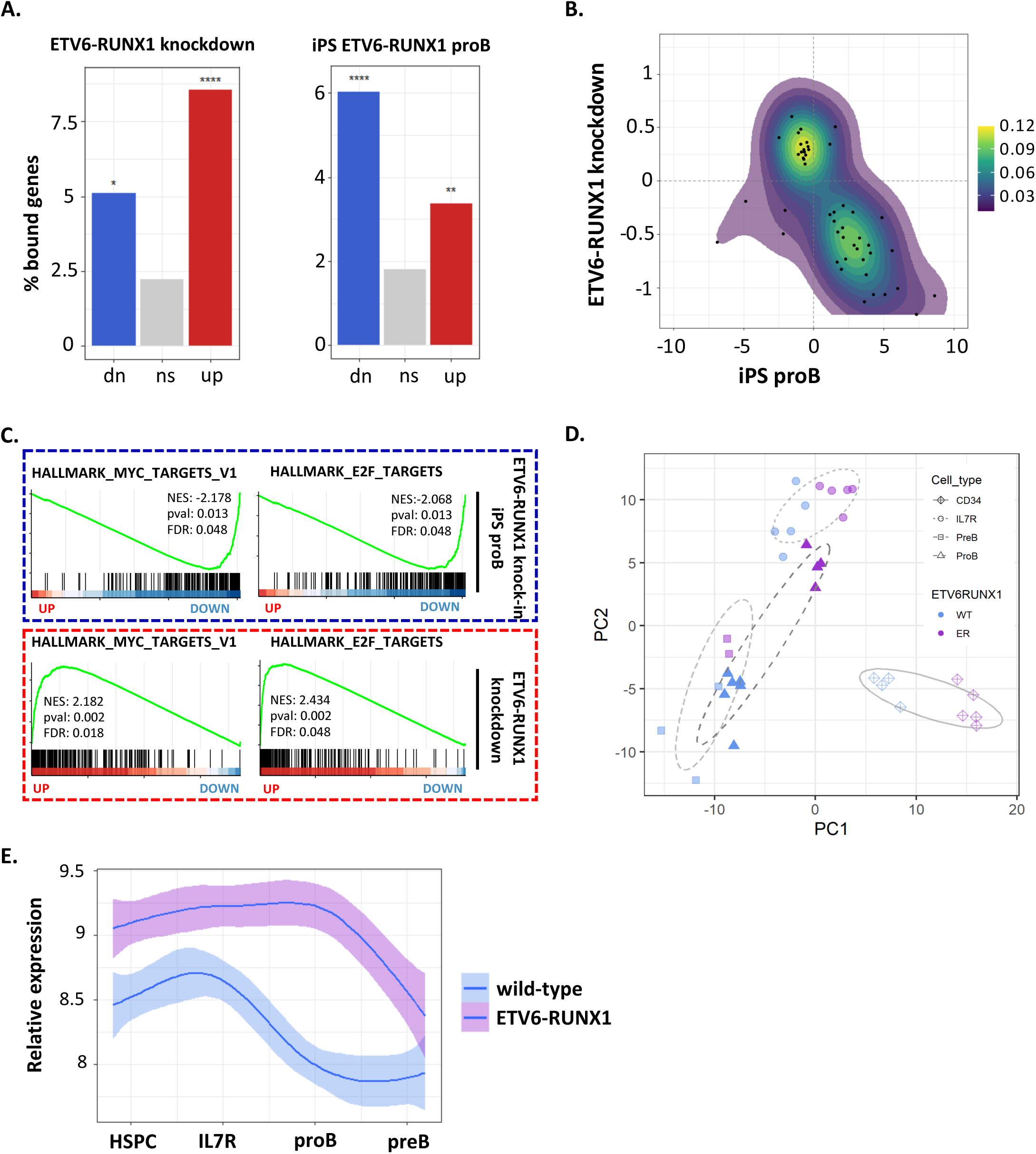
ETV6-RUNX1 induces transcriptional changes indicative of cell cycle repression in pre-leukemia. (A) Bar plot showing percentage of genes associated with a RUNX1 binding event for genes significantly up- or down (dn)-regulated or not significant (ns, p>0.05) upon ETV6-RUNX1 knock down in Reh or ETV6-RUNX1 expression in proB cells derived from IPSCs. **** p<0.0001, *** p<0.001, ** p<0.01, * p<0.05, hypergeometric test. (B) Dotplot with density contours showing log fold change for ETV6-RUNX1 vs wild-type iPSC-derived proB cells and ETV6-RUNX1 knockdown in Reh. Genes displayed have padj<0.1 in both datasets. (C) GSEA for ranked gene lists from ETV6-RUNX1 vs wild-type iPSC-derived proB cells and ETV6-RUNX1 knockdown in Reh. (D) Principal component analysis using label-retaining cell (LRC) signature genes for wild-type and ETV6-RUNX1 expressing iPSC-derived cell populations. (E) Plot showing normalized gene expression (mean +/-95% confidence intervals) in wild-type and ETV6-RUNX1 expressing iPSC-derived cell populations for label-retaining cell leading edge genes derived from GSEA (see Figure S2G).

To identify pathways regulated by ETV6-RUNX1 we used gene set enrichment analysis (GSEA). Comparison to the Hallmark database identified “MYC targets V1”, “E2F targets” and “G2M checkpoint” as significantly enriched in the downregulated genes in ETV6-RUNX1-expressing proB cells (Figure 2C, upper panel) (Subramanian et al., 2005, Liberzon et al., 2015). Conversely, these gene sets were upregulated following ETV6-RUNX1 knockdown, consistent with repression by ETV6-RUNX1 (Figure 2C, lower panel). Since E2F transcription factors are regulators of the G1 to S phase transition and MYC is broadly implicated in cell cycle regulation, these results suggest that ETV6-RUNX1 has a negative impact on the cell cycle of proB cells in a pre-leukemic setting.

While ETV6-RUNX1^+^ ALL invariably has an immature B-cell phenotype, various experimental systems have reported an impact of the fusion on other compartments of the haematopoietic hierarchy (Morrow et al., 2004, Ford et al., 2009). We therefore examined the transcriptional profiles of ETV6-RUNX1-expressing CD34^+^ and IL7R^+^ progenitors, found upstream of proB cells, and of pre-B cells (Boiers et al., 2018). Intriguingly, MYC- and cell cycle-related gene signatures were similarly perturbed in the CD34^+^ and IL7R^+^ progenitors (Figure S2D). However, in the preB population MYC, but not cell cycle signatures, were affected, suggesting that cells escaping the differentiation block do not exhibit a cell cycle impairment (Figure S2D).

Interestingly, even in overt leukaemia a subset of cells remain slow-cycling and are associated with resistance to therapy (Lutz et al., 2013, Ebinger et al., 2016). We examined the impact of ETV6-RUNX1 on a gene signature associated with dormant, label-retaining cells (LRCs) in B-ALL (Ebinger et al., 2016). Principal component analysis (PCA) limited to the LRC signature genes separated samples according to ETV6-RUNX1 status. The greatest impact was on ETV6-RUNX1 expressing proB cells which clustered with the IL7R^+^ population showing that, with respect to the LRC signature, ETV6-RUNX1^+^ proB cells are similar to more immature progenitors (Figures 2D and S2E). In wild-type cells the LRC signature was more highly expressed in HSPC/IL7R+ populations, becoming downregulated in proB cells. Notably, this downregulation was delayed in ETV6-RUNX1^+^ cells until the preB stage (Figures 2E). These results imply that the fusion causes a cell cycle impairment in early progenitors that appears to resolve in the more mature compartments.

In summary, we demonstrate that ETV6-RUNX1 acts predominantly as a transcriptional repressor, both as a first-hit and in the context of full blown leukemia with acquired second hits. In addition we show that ETV6-RUNX1 represses cell cycle-related signatures and that proB cells, corresponding to the point of differentiation arrest in precursor B-ALL, are most significantly impacted.

### Mass cytometry reveals an accumulation of ETV6-RUNX1 expressing progenitor cells in S phase

Our transcriptional analysis strongly suggests that ETV6-RUNX1 has an impact on the cell cycle of B-cell progenitors. Given the limited numbers of differentiated cells obtained with the hIPSC ETV6-RUNX1 model, to directly assess whether cell cycle is altered upon ETV6-RUNX1 expression, we used mass <u>cy</u>tometry <u>t</u>ime-<u>o</u>f-flight (CyTOF) which enables the simultaneous interrogation of surface and intracellular markers across multiple barcoded samples (Table S2, see Methods) (Behbehani et al., 2012). Two *ETV6-RUNX1* knock-in hiPSC clones and two control lines, the parental MIFF3 and an ETV6-RUNX1 clone that has been reverted to wild-type, were subjected to a 30-31 day differentiation protocol that results in a mixture of immature B-cells, progenitors and other haematopoietic cell types (Figure S2C) (Boiers et al., 2018, Carpenter et al., 2011). Viable, CD45^+^ single cells were clustered and visualized with SOM (Self-organizing map) and UMAP (Uniform Manifold Approximation and Projection) dimensionality reduction (see Methods). Clusters were manually annotated according to surface marker expression levels: CD45 (CD45^+^CD45RA^-^CD34^-^CD19^-^), CD45RA (CD45^+^CD45RA^+^CD34^-^CD19^-^), HSPCs (CD45^+^CD34^+^CD19^-^), CD19-low (CD45^+^CD34^lo^CD19^lo^) and preB (CD45^+^CD34^-^CD19^+^) cells (Figure S3A). Consistent with our previous work, our analysis showed that both fusion-expressing lines have markedly decreased B-cell output while an increased proportion of HSPCs and CD19-low cells was observed, indicative of a partial block in B-cell differentiation (Figures 3A and 3B).

**Figure 3.**
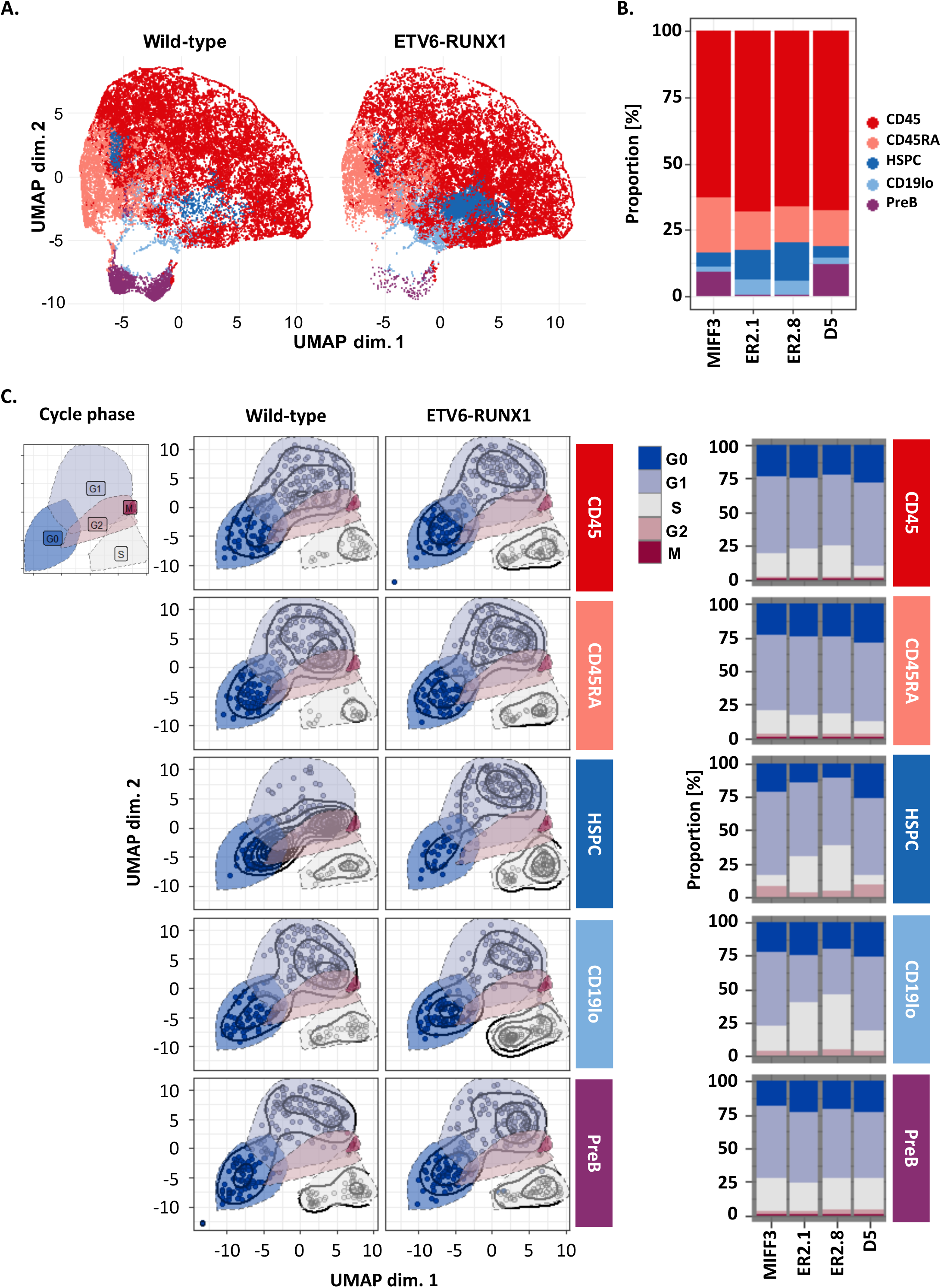
Mass cytometry reveals an accumulation of *ETV6-RUNX1* expressing progenitor cells in S phase. (A) Dotplots of UMAP dimensionality reduction for cell surface markers CD45, CD45RA, CD34, and CD19. Colours represent manually annotated SOM clusters. 10000 cells are shown in each plot. See also Figure S3A. (B) Barplot showing proportion of cells in each population annotated in (A) (C) Dotplots with density contours for UMAP dimensionality reduction based on cell cycle markers pRb, IdU, CycB1 and pHisH3, plotted separately for each of the populations indicated in (A). Populations were randomly downsampled to display equal numbers of cells in each plot. Barplots show proportion of cells in each phase of the cell cycle. Colours represent manually annotated cycle phases based on SOM clusters. See also Figure S3B.

To assess the cell cycle status of the different, immuno-phenotypically defined cell populations, we used clustering to assign each cell to a cell cycle phase based on the markers pRb, IdU, CycB1 and pHisH3 (Figure S3B) (Behbehani et al., 2012). Strikingly, ETV6-RUNX1-expressing HSPC and CD19-low populations had an altered cell cycle profile with an increased proportion of cells in S-phase as compared to controls, suggesting an impaired progression through S-phase (Figure 3C). This difference was not observed in preB cells consistent with our transcriptional analysis suggesting that cells escaping the differentiation block resolve the cell cycle impairment (Figure S2D).

Taken together, our data show, in a developmentally relevant “first-hit” pre-leukemic model, that ETV6-RUNX1 alters the cell cycle profile of early compartments within the B cell differentiation hierarchy. It is paradoxical that a leukemic oncogene should impair cell cycle. We therefore explored its relationship to native RUNX1, a known regulator of cell cycle (Friedman, 2009).

### ETV6-RUNX1 binds DNA through the Runt domain, competing with native RUNX1

Our data suggest that ETV6-RUNX1 hijacks the native RUNX1-driven network by binding to RUNX1 targets through the runt domain. Exploring this in an ETV6-RUNX1^+^ context is difficult, in that a pan-RUNX1 antibody will recognize both the fusion protein and native RUNX1. To directly interrogate their relationship, we used the t(5;12) NALM-6 pre-B ALL cell line (carrying ETV6-PDGFRB fusion, Table S4) which expresses wild-type RUNX1, generating a derivative expressing V5-tagged ETV6-RUNX1, facilitating independent immunoprecipitation of RUNX1 and the fusion. Further, to directly assess the requirement for the Runt and helix-loop-helix (HLH) pointed domains for ETV6-RUNX1 function, we expressed two mutant versions of the fusion carrying a loss-of-function point mutation in the runt domain (R139G) and a HLH deletion (ΔHLH) respectively (Figure S4A).

Comparison of RUNX1 and V5-*ETV6-RUNX1* ChIP-seq datasets revealed a strong correlation of the binding affinities of the two transcription factors, consistent with their having similar DNA-binding properties (Figure 4A, 4B, red dashed line, S4B, S4C and S4D). The most significantly enriched motifs across all binding sites were RUNX and ETS as observed for native ETV6-RUNX1 ChIP in Reh and patient samples (Figures 4C, 1A and Table S3).

**Figure 4.**
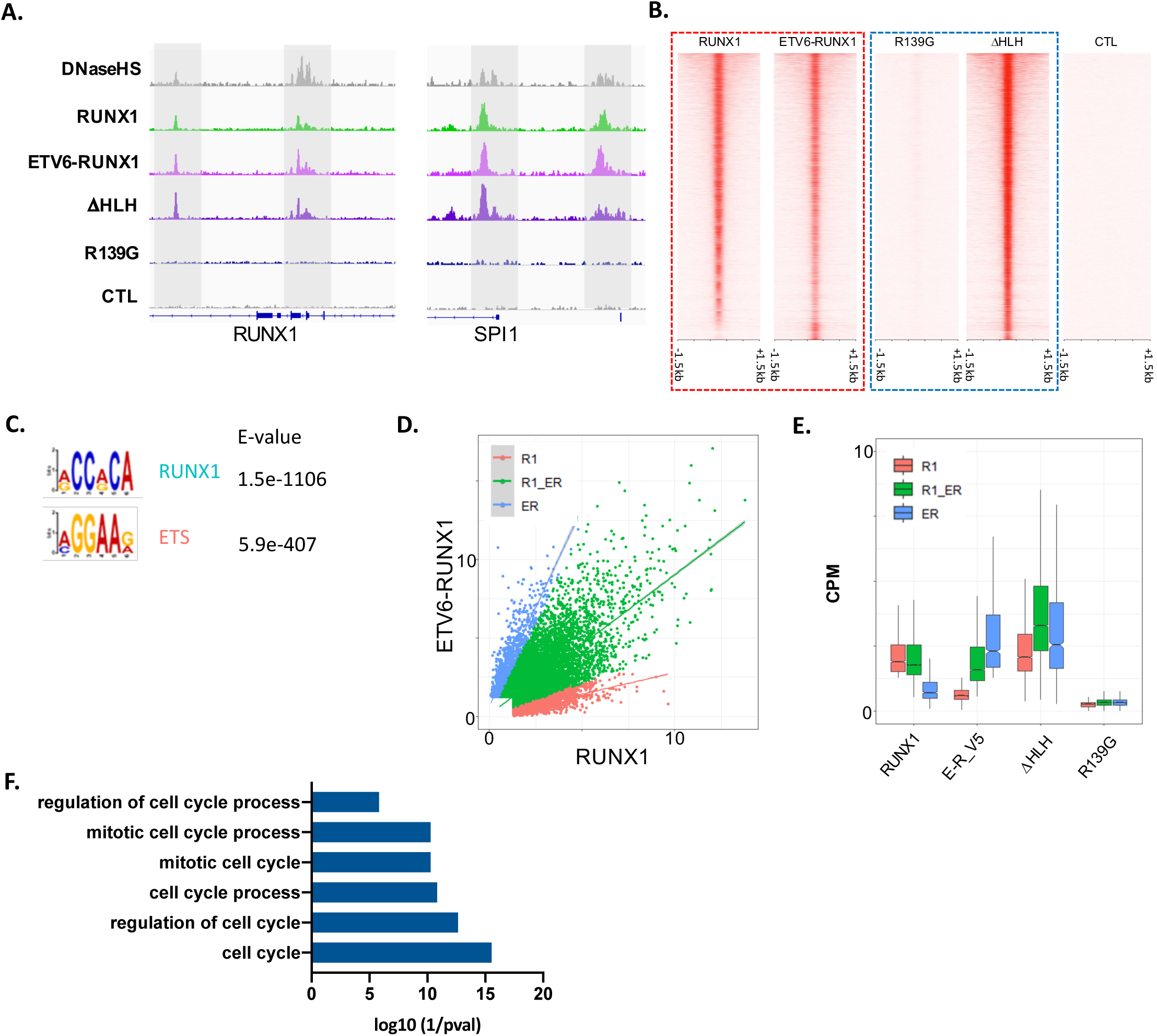
ETV6-RUNX1 binds DNA through the Runt domain, competing with native RUNX1. (A) Integrative Genomics Viewer (IGV) screenshots of RUNX1 and ETV6-RUNX1 binding and DNase1 hypersensitivity (DNaseHS) in NALM6 at the *RUNX1* and *SPI1* loci. R139G: loss-of-function point mutation in the Runt domain of ETV6-RUNX1. ΔHLH: deletion of helix-loop-helix pointed domain of ETV6-RUNX1. CTL: V5 ChIP in NALM6 lacking V5-tagged protein. (B) Heatmaps showing ChIP-seq signal across all identified RUNX1/ETV6-RUNX1 binding sites in a 3kb window centred on peak summits. (C) MEME enrichment of motifs in sites bound by RUNX1 and/or ETV6-RUNX1. (D) Dotplot of CPM for RUNX1 vs ETV6-RUNX1 ChIP. Colours show peaks classified as more highly bound by RUNX1 (R1), ETV6-RUNX1 (ER) or similarly bound by both (R1_ER). (E) Boxplot of CPM for peaks classified in (D) for RUNX1 or ETV6-RUNX1 ChIP datasets. (F) Bar plot showing enrichment of GO-terms relating to cell cycle in genes mapped to R1_ER peaks as classified in (D).

ChIP-seq in the R139G mutant line showed an almost complete loss of binding, formally proving that DNA interaction is dependent on the Runt domain (Figures 4A, 4B, blue dashed line, S4B, S4C and S4D). The fusion inherits the HLH domain from ETV6, which mediates dimerization between members of the ETS family of transcription factors. Interestingly, the ΔHLH exhibited increased binding affinity, suggesting that the HLH domain may in fact inhibit interaction with DNA (Figures 4B, blue dashed line, S4B, S4C and S4D). To explore this further we categorised ChIP peaks into those more strongly bound by the fusion (ER) or RUNX1 (R1) or similarly bound by both (R1_ER) (Figures 4D and 4E). ER peaks tended to be further from the TSS and a slightly higher proportion had one or more RUNX motifs, reflecting the increase in RUNX1 motif frequency with distance from the TSS (Figures S4E and S4F). All three categories of peaks were strongly bound by the HLH mutant suggesting that the pointed domain is responsible for the diminished binding affinity of the fusion protein at R1 peaks (Figure 4E). Taken together these data confirm that ETV6-RUNX1 is able to bind to the same target sites as RUNX1 through the Runt domain, although its distribution across those peaks is altered by the HLH domain.

As a first-hit ETV6-RUNX1 is expressed together with wild-type RUNX1 and the two proteins must compete for binding and regulation of target genes. In the NALM-6 model genes associated with sites bound by both RUNX1 and the fusion (R1_ER) are enriched in GO-terms relating to the cell cycle suggesting that the cell cycle phenotype we observe in our pre-leukaemia model results from disruption of the RUNX1-regulated transcriptome (Figure 4F).

### Defining a core RUNX1 program in B-ALL reveals antagonism between ETV6-RUNX1 and native RUNX1 in cell cycle regulation

Since RUNX1 has been shown to regulate cell cycle progression, we hypothesized that the cell cycle phenotype induced by ETV6-RUNX1 as a first-hit is largely due to dysregulation of the RUNX1 program (Friedman, 2009). We adopted an integrative approach to define a core RUNX1-driven transcriptional network in this disease and extended our analysis from NALM-6 to include RUNX1 ChIP-seq from two additional cell lines representing major B-ALL subtypes.

RUNX1 binding was highly correlated across the three cell lines with 10178 common sites identified, enriched for RUNX and ETS motifs, and mapping to 6843 genes (Figure S5A and S5B). GO term analysis of shared targets identified “cell cycle” as the most enriched biological process, suggesting it is a core RUNX1 function across B-ALL subtypes (Figure S5C).

While childhood and adult B-ALL share some molecular features, they are distinct in their development, prognosis and outcome. To assess the extent to which RUNX1 regulome is conserved across B-ALL subtypes associated with distinct age groups, we examined the transcriptional program engaged upon RUNX1 inactivation by depleting RUNX1 in five cell lines, representing a range of driver mutations found in both childhood and adult ALL, namely ETV6-RUNX1 (Reh), MLL-AF4 (RS4;11), E2A-PBX1 (RCH-ACV), ETV6-PDGFRB (NALM-6) and BCR-ABL (TOM-1) (Table S4), all of which express native RUNX1. To minimize off-target effects and identify high confidence targets we used multiple shRNAs against RUNX1 and performed RNA-seq on biological replicates for each condition (Figures 5A, S5D, red asterisks, and S5E). DEGs were defined for individual cell lines (see Methods) and overlapped to reveal a core set of 321 common DEGs. Of these 247 were up- and 74 downregulated, enriched in GO terms relating to apoptosis and cell cycle/DNA replication respectively (Figures 5B and 5C). We next ranked genes according to their response to RUNX1 knockdown across all five cell lines. The most highly differential genes had a significantly greater association with RUNX1 binding, regardless of expression directionality, indicating that RUNX1 can act as both repressor and activator (Figure S5F). GSEA revealed p53 among the most significantly upregulated pathways, while E2F and MYC targets were significantly enriched in downregulated genes (Figure 5D). This is consistent with wild-type RUNX1 positively regulating cell cycle progression, in direct contrast to ETV6-RUNX1 (Figures 5E, 2 and 3), and suggests that in t(12;21) B-ALL the fusion and native RUNX1 compete with one another with opposing effects on the expression of cell cycle-associated genes.

**Figure 5.**
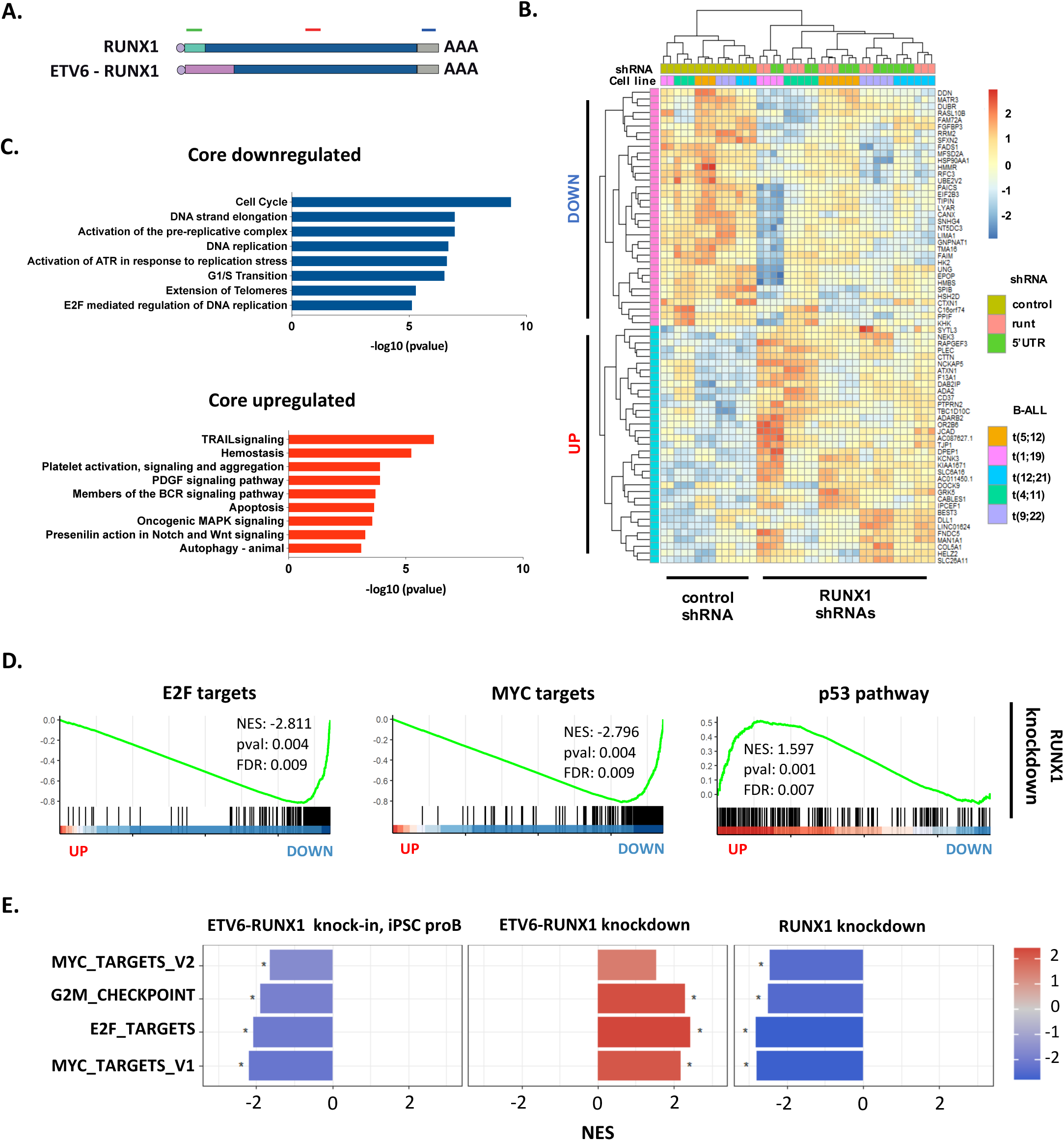
Defining a core RUNX1 program in B-ALL reveals antagonism between ETV6-RUNX1 and native RUNX1 in cell cycle regulation. (A) Schematic showing shRNAs targeting RUNX1 and/or ETV6-RUNX1. (B) Heatmap showing relative expression of the 25 most significantly up- or down-regulated genes following RUNX1 knockdown across five B-ALL cell lines. (C) Bar plots showing significantly enriched GO-terms in the “core” up- or down-regulated genes. (D) Plots of GSEA results for the indicated gene sets against a list of genes ranked from the most significantly up- to the most significantly down-regulated following RUNX1 knockdown. (E) Bar plot showing normalized enrichment scores (NES) for GSEA analysis of the indicated gene sets against ranked lists from ETV6-RUNX1 knock-in iPSC-derived proB cells, ETV6-RUNX1 knockdown and RUNX1 knockdown. * padj < 0.05.

### A balance between CBF complex and ETV6-RUNX1 acting on the P53 regulated cell cycle-apoptosis axis promotes a silent pre-leukemic state

To better understand the apparent competition between RUNX1 and ETV6-RUNX1 in cell cycle regulation, we made use of a meta-analysis categorizing genes according to the number of studies in which a gene is implicated in the cell cycle (Fischer et al., 2016). Higher scoring genes were associated with increased RUNX1 binding and reduced expression upon RUNX1 knockdown showing that cell cycle genes are enriched for direct targets of, and positively regulated by, RUNX1 (Figure 6A and 6B). In contrast, ETV6-RUNX1 negatively regulates the same target genes, clearly demonstrating its opposition to RUNX1 with respect to cell cycle regulation (Figure 6B).

**Figure 6.**
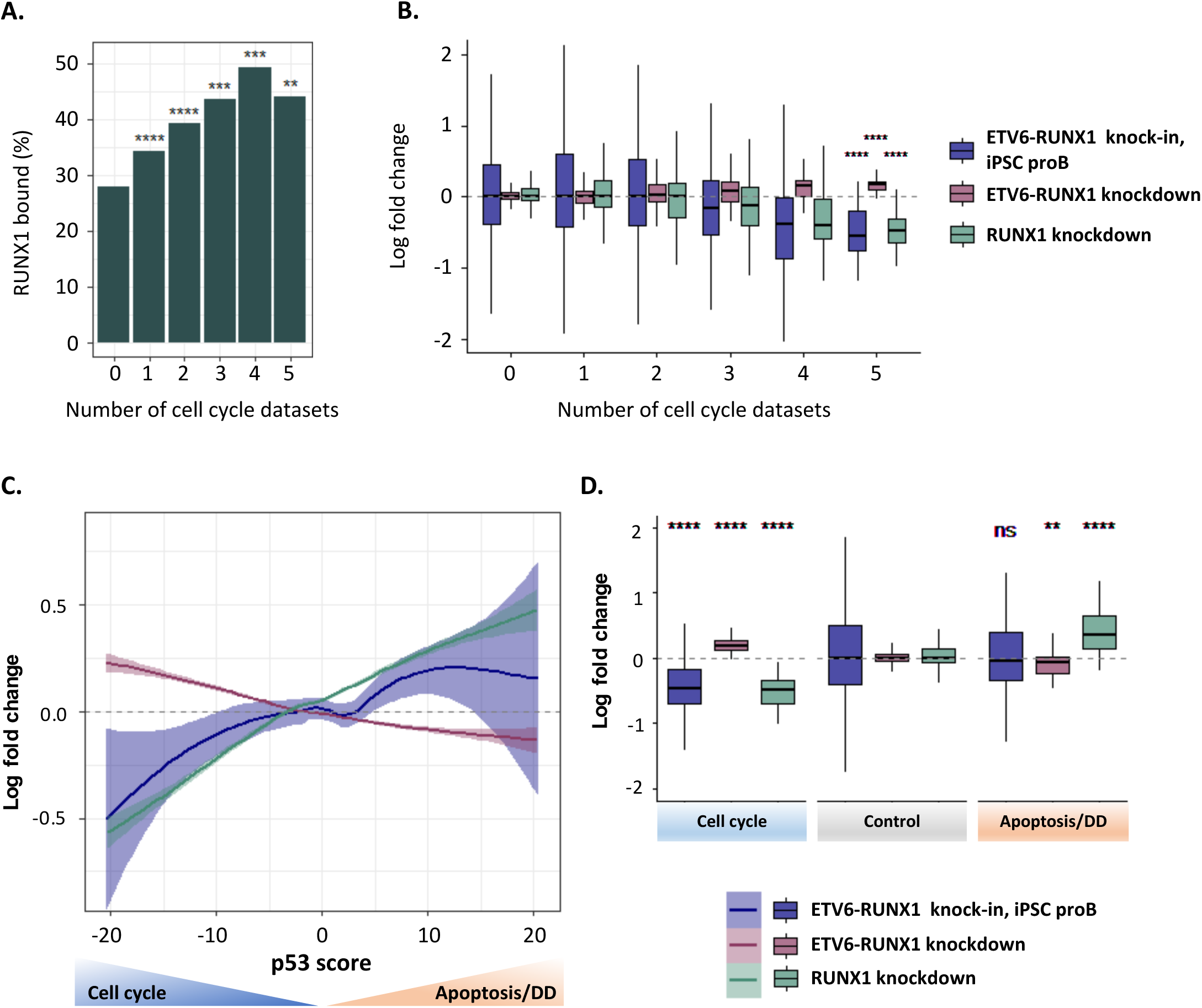
A balance between CBF complex and ETV6-RUNX1 on the P53 regulated cell cycle-apoptosis axis promotes a silent pre-leukemic state. (A, B) Comparison of RUNX1 and ETV6-RUNX1 regulomes to cell cycle meta-analysis (Fischer et al., 2016). Genes were binned according to the number of data sets in which they were defined as cell cycle genes. Bar plot shows the percentage of genes associated with a RUNX1 binding event (A) while boxplots show log fold change following RUNX1 or ETV6-RUNX1 knockdown or in ETV6-RUNX1 knock-in iPSC-derived proB cells (B). **** p<0.0001, *** p<0.001, ** p<0.01, * p<0.05, ns = not significant, unpaired t-test. (C) Plot of mean log fold change for genes binned according to their p53 score for ETV6-RUNX1-expressing ProB cells, ETV6-RUNX1 knockdown, RUNX1 knockdown, and CBFi (AI-14- 91) treatment. Negative p53 scores are associated with cell cycle genes while positive scores are associated with direct p53 targets involved in apoptosis and DNA damage (DD) response (Fischer et al, 2016). See Figure S8A. (D) Boxplot of log fold change for genes grouped into cell cycle (p53 score <= −17), apoptosis/DNA damage (DD) (p53 score >= 17) or control (−17 < p53 score < 17) for ETV6-RUNX1-expressing ProB cells, ETV6-RUNX1 knockdown, RUNX1 knockdown, and CBFi (AI-14-91) treatment. ns = not significant, **** p<0.0001, *** p<0.001, ** p<0.01, * p<0.05, unpaired t-test.

To establish whether there is an association with a particular cell cycle phase, we next analyzed targets of the DREAM complex, RB/E2F, and FOXM1 (Fischer et al., 2016). All three sets were significantly overrepresented among genes negatively regulated by ETV6-RUNX1 while DREAM and RB/E2F were significantly enriched in RUNX1-activated genes (Figure S6A). The strong association with DREAM and RB/E2F implies opposing roles in the regulation of quiescence and G1/S progression and is in agreement with the GSEA results highlighting “E2F targets” as counter-regulated by RUNX1 and ETV6-RUNX1 and with our first-hit iPSC model, where the most marked impact of ETV6-RUNX1 is observed in S-phase (Figures 5A and 3).

Notably, RUNX1 knockdown leads to upregulation of genes associated with p53 activation and apoptosis but this is not observed in our ETV6-RUNX1 knock-in (Figures 5C, 5D and 2). To further explore this difference in behavior, we made use of a p53 meta-analysis which ranks genes according to the number of studies in which they are identified as responding to p53 activation and which broadly divides the p53 regulome into two classes: i) p53 downregulated genes, associated with cell cycle regulation; and ii) p53 upregulated genes, enriched for direct transcriptional targets of p53, associated with DNA damage and induction of apoptosis (Fischer et al., 2016). We compared the ETV6-RUNX1- and RUNX1-regulated transcriptomes to the p53 regulome. All datasets displayed high correlation, with a definite inverse correlation between ETV6-RUNX1 and CBF complex activities (Figure 6C). However, while the p53-downregulated, cell cycle class was overrepresented in both CBF-activated and ETV6-RUNX1-repressed gene sets, the apoptotic class was found to be significantly enriched only in genes upregulated upon RUNX1 inactivation (Figures 6D and S6B).

These data show that a fine balance exists between ETV6-RUNX1 and the native RUNX1. The fusion alters cell cycle without inducing apoptosis, establishing a clinically covert pre-leukemic state whereas direct depletion of RUNX1 impacts cell cycle and in addition induces signatures of apoptosis. This was observed in five B-ALL subtypes and suggests that dependence on RUNX1 activity is a general feature of B-ALL (Figures 5 and S5).

### B-ALL cell lines and primary patient cells are dependent on RUNX1 activity for survival *in vitro* and *in vivo*

RUNX1 knockdown leads to upregulation of genes associated with p53 activation and apoptosis, suggesting a dependence on RUNX1 not only for proliferation, but for survival of B-ALL cells.

To functionally assess the requirement for RUNX1 in leukemia, we depleted RUNX1 *in vitro* in four B-ALL cell lines and a control chronic myeloid leukemia (CML) cell line and monitored proliferation of leukemic cells over nine days (Figures S5D, red asterisks, and 7A, Table S4). B-ALL cells transduced with any of the RUNX1 targeting shRNAs showed greatly reduced proliferative capacity compared to non-targeting control, while the t(9;22) CML cell line K562 was unaffected, consistent with previous reports (Goyama et al., 2013). While differences in sensitivity between shRNAs reflected the degree of RUNX1 knockdown, we noted that even small perturbations of RUNX1 levels caused proliferation arrest (Figures S5D, 5’UTR shRNA and 7A).

We next subjected RUNX1 depleted cells to mass cytometry analysis to assess cell cycle distribution and apoptosis in these cells. We noted increase in G0 and/or G1-phases and an increase in cleaved Caspase-3 and Cisplatin-positive populations in RUNX1 depleted cells, demonstrating that diminished RUNX1 activity causes a delay in cell cycle progression, as well as an increase in apoptosis (Figures 7B and S7A).

**Figure 7.**
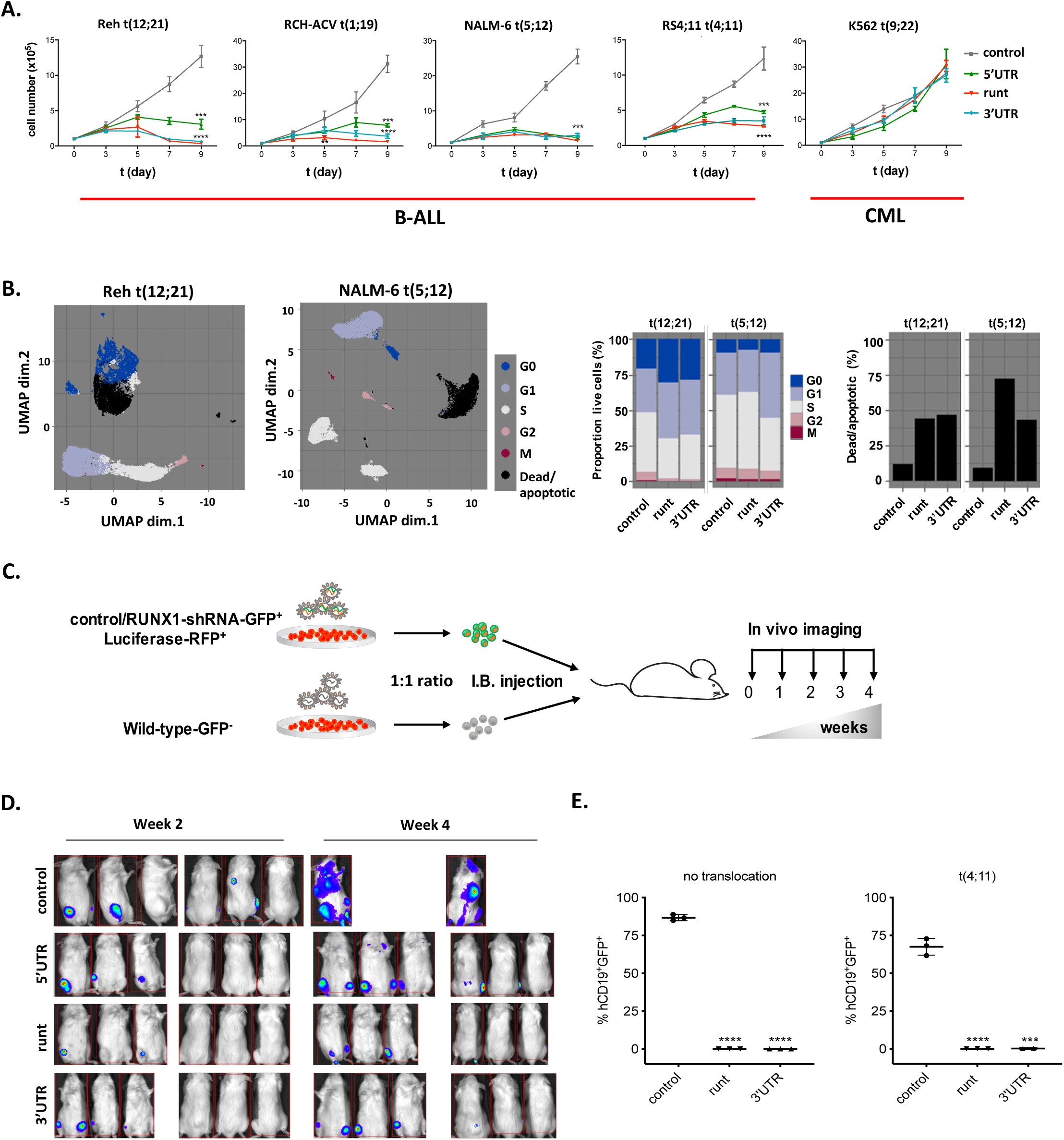
B-ALL cell lines and primary patient cells are dependent on RUNX1 activity for survival *in vitro* and *in vivo*. (A) Growth curves for the indicated cell lines over 9 days following transduction with shRNAs targeting RUNX1 (Figure 5A) or a non-targeting control. (B) Mass cytometry analysis for the indicated cell lines following transduction with shRNAs targeting RUNX1 (Figure 5A) or a non-targeting control. UMAP projections based on cell cycle and viability/apoptosis markers. Colours represent manually annotated SOM clusters. Barplots show proportion of live cells in the indicated cell cycle phases and the proportion of dead/apoptotic cells. (C) Schematic of competitive engraftment experiments. NALM-6 cells transduced with shRNAs (GFP^+^) and stably expressing a luciferase/RFP reporter were mixed 1:1 with non-transduced (GFP^-^) cells and injected into NSG mice. Mice were imaged weekly for luciferase activity and culled for end-point analysis (4 weeks) of leukaemic engraftment. (D) Bioluminescence imaging at 2 and 4 weeks (end point) after injection of samples transduced with shRNAs targeting RUNX1 or control shRNA (see Figure 7C). (E) Plots showing end point analysis of leukaemic engraftment of two patient samples with indicated cytogenetics transduced with non-targeting or RUNX1-shRNAs. Human (CD19^+^), gated on human CD45^+^ cells, were assessed for the proportion of shRNA-transduced (GFP^+^) vs wild-type GFP^-^ cells (Figure S7C). **** p<0.0001, *** p<0.001, unpaired t-test.

To directly assess the impact of RUNX1 depletion on the engraftment and proliferative capacity of B-ALL cells in vivo we used bioluminescence imaging to follow leukaemic propagation of RUNX1 shRNA or control shRNA-transduced cells (1:1 ratio) in a competitive setting (Figure 7C). NOD SCID gamma (NSG) mice injected intra-tibia with NALM-6 cells (see Methods) showed initial engraftment of all samples at the injection site but while control shRNA-transduced cells were observed at secondary sites, such as the spleen and brain, RUNX1-depleted cells failed to disseminate (Figures 7D, S7B and S7C). In addition, splenic size was visibly decreased in RUNX1-depleted mice, consistent with a diminished leukemic burden in those animals (Figure S7D).

To determine if primary, patient-derived B-ALL cells are similarly dependent on RUNX1 in vivo, we transduced two clinically aggressive pediatric B-ALL patient samples with the two most effective RUNX1 shRNAs and engrafted them into NSG mice (Table S5, see Methods). FACS analysis of total bone marrow at the end point (3 months) showed that shRNA-RUNX1 cells are outcompeted by the non-transduced control cells and were barely detectable by FACS (Figure 7E and S7E), while in animals engrafted with control shRNA-transduced cells both GFP^+^ and GFP^-^ populations were present.

In sum, our results clearly demonstrate dependence on RUNX1 for survival across B-ALL, suggesting it may serve as a therapeutic target.

### An allosteric CBFβ inhibitor mimics RUNX1 depletion phenotype and offers a targeted treatment for B-ALL

RUNX1 is part of the CBF complex and its binding to DNA is stabilized by the CBFβ subunit. We therefore assessed if disruption of their interaction would phenocopy the effects of loss of RUNX1. We first depleted CBFβ using an shRNA (Figure S5D, blue asterisk) and compared transcriptional changes to RUNX1 knockdown across the five cell lines described above (Table S4). For significantly dysregulated genes the direction of change was highly correlated, demonstrating that depletion of CBFβ results in reduced RUNX1 transcriptional activity (Figure S8A). Similarly to RUNX1 depletion, CBFβ knockdown resulted in reduced growth in vitro of four B-ALL cell lines, while K562 cells were unaffected (Figure 8A). Strikingly, CBFβ-depleted NALM-6 cells and two primary B-ALL patient samples failed to engraft and proliferate in vivo, recapitulating RUNX1 dependence for survival (Figures 8B, 8C, S8B and S8C).

**Figure 8.**
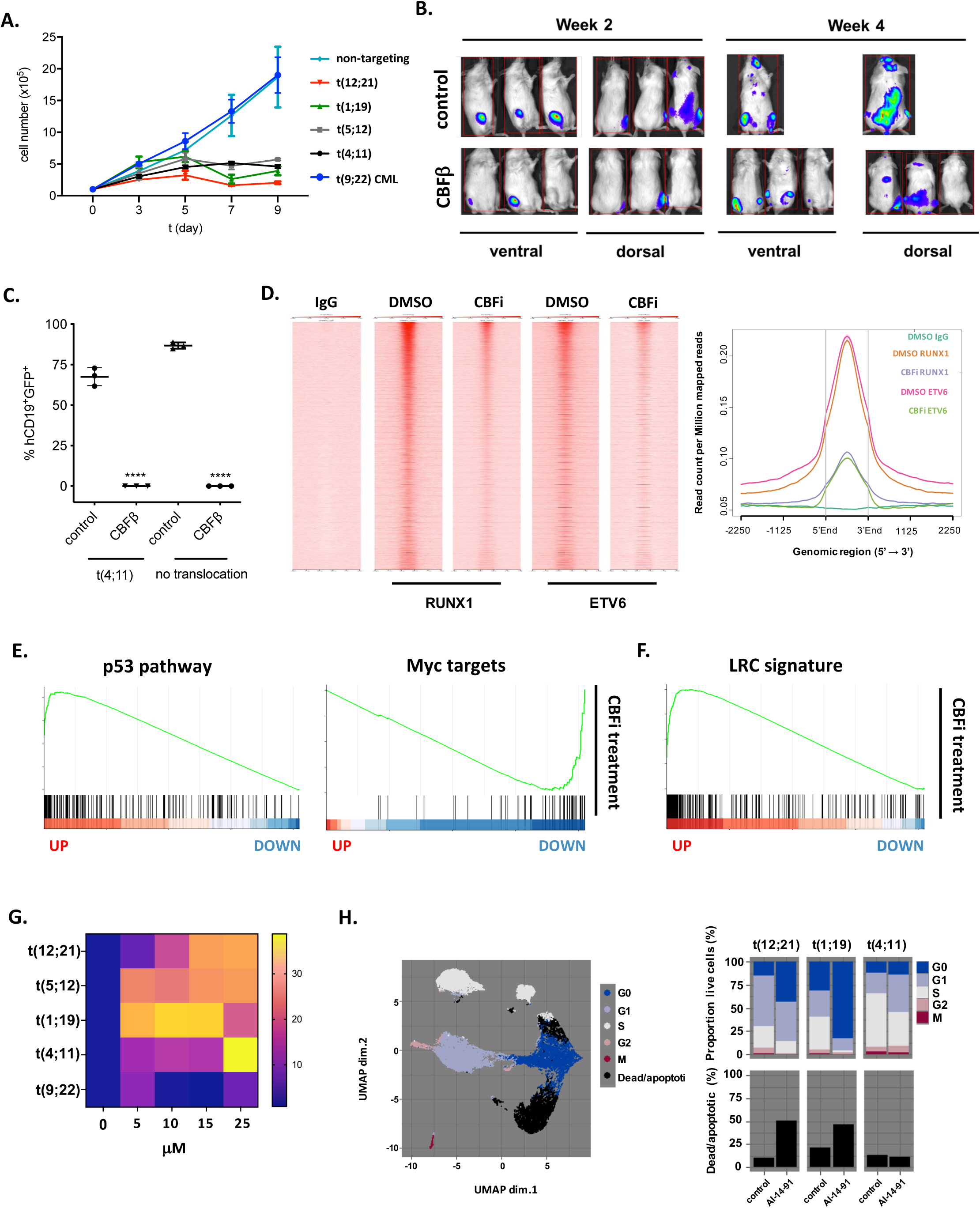
An allosteric CBFβ inhibitor mimics RUNX1 depletion phenotype and offers a targeted treatment for B-ALL. (A) Growth curves for the indicated cell lines over 9 days following transduction with an shRNA targeting CBFβ or a non-targeting control. Error bars indicate standard deviation (n=3). (B) In vivo bioluminescence imaging of NOD/SCID mice competitively engrafted with shGFP^+^Luc^+^ and wild-type GFP^-^Luc^-^ NALM-6 cells (see methods and Figure 7C). Ventral and dorsal views of mice at weeks 2 and 4 (end point); control (non-targeting shRNA); CBFβ-CBFβ-targeting shRNA. (C) Summary endpoint FACS analysis of leukaemic engraftment of two patient samples with indicated cytogenetics transduced with a non-targeting or CBFβ shRNA. Human (CD19^+^), gated on human CD45^+^ cells were assessed for the proportion of shRNA-transduced (GFP^+^) vs wild-type GFP^-^ cells (Figure S7C). **** p<0.0001, unpaired t-test. (D) Heatmap (left) and average profiles (right) of ChIP-seq signal across all RUNX1 and ETV6-RUNX1 binding sites in a 3kb window centred on peak summits from control (DMSO) or CBFi (AI-14-91) treated Reh (ETV6-RUNX1^+^ ALL) following immunoprecipitation with RUNX1, ETV6 or control (IgG) antibodies. (E) GSEA plots for p53 and Myc against genes ranked from most significantly up-to most significantly down-regulated following CBFβ knockdown. (F) GSEA plot of LRC signature against genes ranked from most significantly up-to most significantly down-regulated following CBFβ knockdown (G) Heatmap showing cell death in response to AI-14-91 (CBFi). The indicated cell lines were cultured with the increasing concentrations of CBFi for 72h. Values are relative Celltox green signal normalised to untreated (0μM) cells. (H) Mass cytometry analysis for three B-ALL cell lines Reh, t(12;21), RCH-ACV t(1;19) and RS4;11 t(4;11) treated with 15uM CBFi (AI-14-91) for 48h. UMAP projections based on cell cycle and viability/apoptosis markers. Colours represent manually annotated SOM clusters. Barplots show proportion of live cells in the indicated cell cycle phases and the proportion of dead/apoptotic cells.

This genetic proof-of-principle suggested that pharmacological disruption of the CBF complex would be sufficient to induce cell cycle arrest and apoptosis in B-ALL. We therefore tested the effect of AI-14-91 (hereafter referred to as CBFi), an allosteric inhibitor of the CBFβ-RUNX1 interaction (Illendula et al., 2015, Illendula et al., 2016). ChIP-seq analysis of Reh cells treated with CBFi showed reduction of global binding patterns of RUNX1 (Figure 8D). Notably, ETV6-RUNX1 binding was similarly reduced, showing that interaction with DNA remains CBFβ-dependent in the fusion in line with Roudaia et al. (Roudaia et al., 2009). Transcriptional changes following treatment with CBFi were highly correlated to those observed upon RUNX1 knockdown, with a similar impact on pathways relating to Myc and p53 (Figures S8D and 8E). This demonstrates that treatment with CBFi results in inhibition of RUNX1-dependent transcription. In addition, the LRC signature used previously showed a significant enrichment in the CBFi upregulated genes, recapitulating the effect of ETV6-RUNX1 (Figure 8F, 2D and 2E).

To test the functional impact of the inhibitor we treated four B-ALL and a CML cell line with increasing concentrations and measured their survival and proliferation in vitro. Strikingly, all B-ALL cells exhibited a dose-dependent reduction in viability, while t(9;22) CML cells were unaffected, corroborating our genetic data and consistent with a pan-B-ALL dependency on CBF activity (Figure 8G). Similarly, colony forming capacity was significantly reduced in cells treated with CBFi (Figure S8E). Furthermore, CyTOF analysis revealed an increase in cell death, accompanied by increased expression of cleaved Caspase3, and an increase in G0 and/or G1 cells consistent with CBF complex being required for B-ALL survival and cell cycle progression (Figure 8H).

To directly assess the contribution of CBF complex to cell cycle progression we arrested cells in G1 or G2 through pharmacological inhibition of CDK4/6 or CDK1 respectively and monitored re-entry to the cell cycle in the presence or absence of CBFi. We observed delayed progression from G1 through S phase but no impact on cell cycle progression from G2 (data not shown), demonstrating a role for CBF-regulated transcription in G1/S phases of the cell cycle consistent with reduced expression of E2F targets (Figures S8F and 5D).

Finally, we tested the effect of CBFi on CD34^+^ human bone marrow cells in vitro. Of note, no significant reduction in viability was observed in the timeframe measured, in agreement with previous work showing that RUNX1 is dispensable for normal HSPC-function (Figure S8G) (Cai et al., 2011).

Taken together, our data demonstrate that allosteric inhibition of the CBF complex phenocopies genetic depletion of either of the two subunits, opening a novel therapeutic avenue in B-ALL.

## Discussion

As ETV6-RUNX1 arises *in utero* as an initiating lesion, we sought to understand its function as a “first-hit”, exploiting an iPSC model of pre-leukemia. We show that ETV6-RUNX1, through repression of RUNX1 targets, downregulates transcriptional pathways associated with cell cycle progression. RUNX1 has previously been described to be a cell cycle regulator and analogous findings in AML show that the onco-fusion RUNX1-ETO compromises cell cycle and high levels of RUNX1-ETO induce apoptosis (Friedman, 2009, Martinez-Soria et al., 2019, Ptasinska et al., 2014, Mandoli et al., 2016). It is perhaps counterintuitive that an oncogene should oppose cell cycle. However, ETV6-RUNX1 alone is not sufficient to establish leukemia. Rather, by impeding differentiation and cell cycle progression it may generate a pool of progenitors arrested in their development, vulnerable to second hit mutations. Notably, a common second hit is loss of CDKN2A – encoding p16 and p14 from alternative reading frames – which promote cell cycle due to loss of a CDK inhibitor and survival through MDM2-mediated suppression of p53, respectively (Heltemes-Harris et al., 2011, Sulong et al., 2009). The negative effects of ETV6-RUNX1 on cell cycle may create positive selection for loss of p16. Our data parallel with a recent study demonstrating that expression of RUNX1-ETO in an inducible human embryonic stem cell model causes a quiescent phenotype in early myeloid progenitors (Nafria et al., 2020).

Given that the fusion functions by opposing the native CBF complex, it might be expected that loss of function mutations in *RUNX1* would be observed in B-ALL. However, the opposite appears to be the case with retention or amplification of the wild-type allele. We show here that loss of RUNX1 in B-ALL cells results in cell cycle exit and apoptosis revealing that B-ALL cells are RUNX1 addicted, consistent with the requirement for Runx1 for survival of immature B-cells in mice (Niebuhr et al., 2013). *Runx1* heterozygosity is well tolerated in the B-cell lineage although compound haploinsufficiency with *Ebf1* reveals a role in induction of the B cell transcriptional program (Lukin et al., 2011). Our results suggest that loss of the second allele of *RUNX1* would not be tolerated as indeed was the case in a mouse model where ETV6-RUNX1 expression, together with homozygous loss of *Runx1*, resulted in severe anemia and death of the animals (Schindler et al., 2009). We suggest therefore that t(12;21) is only viable in the continued presence of wild-type RUNX1 and may even create selective pressure for additional copies. That ETV6-RUNX1 is tolerated as a first hit is explained by the absence of p53 activation that is observed upon direct inhibition of CBF. A link between ETV6-RUNX1 and p53 has previously been suggested, with forced expression of the fusion inducing MDM2, which represses p53 activity (Kaindl et al., 2014). Although we observed ETV6-RUNX1 binding at the *MDM2* promoter we did not observe any change in its expression.

*TP53* is rarely lost in B-ALL suggesting that CBF inhibition may provide a route to tumor cell killing without the collateral damage associated with systemic p53 activation. Moreover, if RUNX1 is selectively retained, targeting the CBF complex may provide a means to overcome chemotherapy resistance associated with subclonal heterogeneity (Dobson et al., 2020). However, current CBF inhibitors are not sufficiently potent to be used in the clinic. Future efforts should see more potent compounds developed and will explore combination therapies as a possible route to less toxic treatments.

## Acknowledgements

This work was generously funded by grants from the UK: Blood Cancer UK (Grant #16001), Children with Cancer UK (Grant#17-250) and the Medical Research Council UK (Grant#MR/N000838/1). We thank Cure Cancer@UCL and Mrs Sandra Hamilton for the generous support. We thank John Brown for input on the RNA-sequencing experiments, Mathew Robson for help with in vivo animal imaging, Chela James and Javier Herrero for input on bioinformatic analysis and George Morrow from the UCL Cancer Institute Flow Cytometry Facility for help with Mass Cytometry. We further thank Gill May for critical input on the manuscript.

## Author Contributions

J.P.W., E.M.D., R.N. and T.E. conceived the experiments. J.P.W., E.M.D., C.B., J.B.C., S.G., Y.G. performed the experiments. E.M.D and J.P.W. analysed the experiements. J.P.W. performed the bioinformatics analysis. S.E.R. made the ETV6-RUNX1 iPS lines. A.I. and J.H.B. provided the CBFβ inhibitor. J.H.A.M. and H.G.S. performed the ChIP-seq of NALM-6 mutant lines. J.P.W., E.M.D. and T.E. wrote the manuscript.

## Declaration of Interests

The authors declare no competing interests

## Methods

### Lead Contact

Further information and requests for resources and reagents should be directed to and will be fulfilled by the Lead Contact, Tariq Enver.

### Materials Availability

Plasmids used in this study are available upon request from the Lead Contact.

### Data and Code Availability

Data relating to IPSC ETV6-RUNX1 knock-in model: ArrayExpress, E-MTAB-6382. St Judes paediatric B-ALL: PanALL, SJC-DS-1009

### Cell lines

For full details (age and cytogenetics) please refer to Table S4. Reh, NALM-6, RCH-ACV, RS4;11 and TOM-1 were purchased from DSMZ. K562 were a kind gift from Prof Asim Khwaja, UCL Cancer Institute, Department of Hematology, London, UK. 293T cells were purchased from ATCC. All leukemic cells were grown in RPMI 10% FBS at 37°C and 5% CO_2_. 293T cells were grown in DMEM 10% FBS at 37°C and 5% CO_2_. iPSC cells were maintained and differentiated as in (Boiers et al., 2018).

### Primary patient material

Diagnostic childhood B-ALL bone marrow (BM) samples were obtained upon informed consent and approval of relevant research ethics from patients at the Great Ormond Street Hospital (GOSH). Mononuclear cells were isolated by Ficoll gradient centrifugation and frozen at −80° until further use. For full details (age and cytogenetics) please refer to Table S5. Prior to experiment samples were thawed at 37° and transduced with the respective lentiviral constructs (MOI of 10), incubated for 48h and flow cytometry sorted prior to injection.

### Bone marrow reconstitution assay

All *in vivo* experiments were performed in strict accordance with the United Kingdom Home Office regulations. Animals were housed in individually ventilated cages (IVC) under specific pathogen free (SPF) conditions. Primary childhood B-ALL cells were transplanted into 8-12 weeks old sub-lethally irradiated NOD/SCID IL2Rγ^null^ (NSG) mice (males and females, obtained from Dominique Bonnet’s Lab, The Francis Crick Institute) by an intravenous (IV) injection. Sub-lethal irradiation was achieved with a single dose of 375 cGy. Prior to the procedure, mice were administered acid water for a week and Baytril (resuspended at 25.5 mg/kg in the drinking water) for the 2 weeks following it. Each mouse received a total of 1×10^5^ leukemia cells resuspended in 100μl PBS/0.5% FBS.

### *In vivo* bioluminescence imaging

For *in vivo* imaging studies the lentiviral TWR vector (Firefly Luciferase-Red fluorescence protein), kindly donated by Prof Dominique Bonnet, was used (Gandillet et al., 2011). NALM-6 cells were co-transduced with TWR and LLX3.7-shRNA vectors and RFP^+^GFP^+^ double positive cells were FACS sorted and injected intra tibia. IVIS®Lumina was used to image animals. Prior to imaging animals were anaesthetized with 3% isoflurane and D-Luciferin (Caliper Life Science) was injected intraperitoneally (IP). Anterior and posterior images were taken at different exposure times post injection to determine peak signal. Total bioluminescent signal was assessed as a sum of anterior and posterior signals.

### Flow cytometry

In general, for flow cytometry analysis or sorting cells were centrifuged at 300rcf for 5min and washed in 1ml PBS/2%FBS. Staining was done in 100μl volume. A master mix with appropriate antibodies was prepared and added to each sample. Unstained and single stain controls were included for each experiment and Hoechst 33258 (SigmaAldrich) was used as a viability dye. Prior to staining cells obtained from animals were lysed with 1ml of Red Blood Cell (RBC) lysis buffer (SigmaAldrich) and incubated for 5-10 min at room temperature.

### Cell Cycle Assay

Ki67 staining was used to assess cell cycle status. Cells were washed with 1ml of PBS/2%FBS, resuspended in PBS/2%FBS and fixed with freshly made 3.2%PFA (paraformaldehyde)/PBS for 10min at room temperature in the dark. After washing cells were permeabilized in 300μl cold 90%Methanol/PBS on ice for 30min and then washed twice as before. 25μl of Ki67 antibody was added and samples were incubated at RT for an hour. DAPI (0.5μg/ml) staining was done for 40 min on ice. Cells were washed, resuspended in 300μl PBS/2%FBS and FACS analysed on Gallios® (Beckman Coulter) or ARIAIII (BD). Unstained, isotype and single stain controls underwent the same procedure without addition of the respective antibodies.

### Apoptosis Assay

AnnexinV (BD) and Hoechst 33258 dye were used to detect apoptotic cells. Cells were stained following manufacturer’s protocol, resuspended in 200μl 1xBinding buffer (BD) containing Hoechst58 (1:10000 dilution) and analysed on the Gallios® (Beckman Coulter).

### Knockdown using shRNAs

Small hairpin RNAs targeting RUNX1, CBFβ and ETV6-RUNX1 were designed using several prediction tools or The RNA Consortium (TRC) website. In general, a sequence of 21nt in length ting with G was chosen where possible. The loop sequence (GGGATCCG) was designed to contain the BamHI (GGATCC) restriction site, not present in the LLX3.7 vector, which was then used for selection of positive clones. Sequences were ordered as primers from Integrated DNA Technologies (IDT) website and resuspended in DNase–free water as a 100μM stock concentration. For oligo annealing 1μl of 100μM forward and reverse primers for each shRNA construct were mixed in a PCR tube with 48μl of IDT Duplex Buffer (Integrated DNA Technologies) and cloned into the LLX3.7 lentiviral vector. For TRIPZ vector shRNA were designed as in (Fellmann et al., 2013) and cloned following manufacturer’s instructions. To induce knockdown doxycycline was added at a final concentration of 0.5μg/ml and cells were collected for western blot analysis 72h later. For RNA-seq cells were collected 7 days post induction with Doxycycline.

### Lentiviral production

Lentiviral particles were packaged using the second generation packaging system and for each lentiviral construct three 175cm^2^ flasks of HEK293T cells were used. HEK293T cells were grown to 70-80% confluency and media was changed before adding the DNA mix. For each lentiviral construct a mix of 4.47μg lentiviral vector, 2.98μg of psPAX2 and 2.98μg of pMD2.G was made in a sterile TE Buffer. Opti-MEM^®^ media was added to the DNA mix. FuGENE^®^6 was used as a transfection reagent. The DNA/media/FuGENE^®^6 mix was incubated at RT for 15-30 min and added dropwise to the flask containing HEK293T cells avoiding contact with the plastic. The cells were then incubated for 48h without changing the media and at that point supernatant was collected from each flask. Further 18ml of DMEM were added to each flask and cells were incubated for another 24h. To collect viral particles, supernatants were filtered through a sterile 0.45μm surfactant-free, non-pyrogenic filter into Oak Ridge centrifuge tubes with a seal cap (Nalgene). The tubes were centrifuged at 50 000g for 3h at 4°C, supernatant was discarded and the pellet was resuspended in 100μl of IMDM (FBS-free, volume for one 175cm^2^ flask), aliquoted into screw-cap tubes and kept at −80° for further use.

### Lentiviral transduction

Cell lines were washed with PBS/2%FBS and resuspended in complete medium at a concentration of 1*10^6-3*10^6/ml. 1ml of cells was added into a well of a 6 well plate and virus was added dropwise (volume of virus depending on MOI). 24h later 1ml of media was added, cells were incubated for further 24h and washed twice with PBS/2%FBS prior to FACS analysis or sorting. Primary patient material was aliquoted into 24- or 48-well plates in 200μl StemSpan™ Serum-Free Expansion Medium (SFEM) (STEMCELL Technologies) supplemented with 20% FCS and the following cytokines: 50ng/ml human SCF, 20ng/ml IL-7, 10ng/ml IL-3, /ml FLT-3.

### In vitro proliferation assay

B-ALL cell lines were transduced with control, RUNX1 or CBFβ shRNAs, FACS sorted 48h later and cultured in 6-well plates in 4ml of media. Cells were counted daily for a period of 9 days.

### AI-14-91 treatment

AI-14-91 was provided as powder and resuspended in DMSO to a stock concentration of 40mM. Prior to experiment respective dilutions were made in media and added to the cells. 0.25% of DMSO was used as control.

### CellTox Green assay

Cells were resuspended in media with CellTox™ Green (Promega) in 96-well plates (100 000 cells/well). Fluorescence intensity was measured at (Ex)485/(Em)520nm on Varioscan™ Lux microplate reader (ThermoFischer Scientific) 72h later.

### Colony Formation Assay

B-ALL cells were treated with increasing concentrations of AI-14-91 for 48h, washed and seeded into 1.5 mL of Methocult H4230 (Stem Cell Technologies) supplemented with 50ng/ml human SCF, 20ng/ml IL-7, 10ng/ml IL-3, /ml FLT-3 and plated into 35 mm non-coated plates (430,588, Corning Incorporated, Corning, NY, USA). Plates were incubated for 10–14 days at 37 °C, 5% CO_2_. Colonies produced were counted and classified.

### Palbociclib treatment

Cells were synchronised by incubation with palbociclib (500nM) for 48h. Cells were then washed with PBS and returned to culture in the presence of AI-14-91 (20uM) for 18h. Cells were incubated with Hoechst 33342 (0.5μg/ml) for 30mins and subjected to FACS analysis. For cell cycle analysis sub-G0 cells were excluded. G0/G1 (2N), G2 (4N) and S-phase (2N < S < 4N) cells were gated based on DNA content and quantified.

### Western Blotting

Cells were pelleted, washed with PBS Buffer and lysed (30 min on ice with occasional vortexing) in RIPA Buffer (Sigma Aldrich) supplemented with 1x Protease Inhibitor Cocktail (Roche). Lysates were cleared by centrifugation at 4°C for 15 min at 14000g and total protein extracts were transferred to a new pre-chilled Eppendorf tube. For nuclear extracts (ETV6-RUNX1) the remaining pellet was resuspended in 50μl RIPA supplemented with 1x Protease Inhibitor Cocktail and sonicated for 3-5 min with Picoruptor^®^ (Diagenode). 10μg of proteins were then separated through precast 4-12% NuPAGE^®^ Bis-Tris gels (Invitrogen) and transferred onto a PVDF membrane (Millipore). Protein membrane was blocked and incubated with primary antibody O/N at 4°C. Following 1h incubation with secondary antibodies, membranes were developed using the ImageQuant LAS 4000 System (GE Healthcare). Antibodies used: RUNX1 (Abcam, ab23980), ETV6: (Sigma, HPA000264), CBFβ (Abcam, ab33516), GAPDH (14C10, Cell Signaling #2128).

### Chromatin Immunoprecipitation

Cells were crosslinked with 1% formaldehyde and sonicated to yield chromatin of 100–500 bp. ChIP was performed by as previously described (May et al., 2013) using antibodies: RUNX1 (Abcam, ab23980), ETV6: (Sigma, HPA000264), V5 (Abcam, ab9116), and nonspecific rabbit IgG (Abcam, ab171870). Libraries were gel-purified, 10ng of DNA was amplified and single end sequenced at 36 bp using Illumina NextSeq500/550 High Output Kitv2.5 (75 cycles). For AI-14-91 ChIP-seq Reh cells were treated with 15μM CBF inhibitor for 48h prior to crosslinking.

### Mass Cytometry

iPS or B-ALL cell lines were incubated with IdU at a final concentration of 50μM for 30min at 37°. Cells were then washed with MaxPar Staining Buffer and incubated in 1ml of MaxPar PBS with Cell-ID Cisplatin (final concentration of 1uM) for 5min at room temperature. Following a wash with 5ml of MaxPar Staining buffer, samples were barcoded following manufacturer’s instructions. Up to 20 barcoded samples were then pooled and the sample was counted to determine the amount of surface antibody necessary. As a rule, 1ul of each antibody was used to stain 3×10^6 cells. Pool was incubated with the surface antibody cocktail (see Table S1 for details of all antibodies used) for 30min at RT with vortexing in between. After washing with MaxPar Staining Buffer cells were incubated with 1x Nuclear Antigen Staining Buffer for 30min, followed by a wash with Nuclear Antigen Staining Perm. Cells were counted prior to addition of a cocktail of intracellular antibodies and incubated at RT for 45min with gentle vortexing in between. Cells were then fixed with 1.6% freshly prepared PFA for 10min at RT. After washing cells were incubated for 1h at RT or overnight at 4°C in 1ml MaxPar Fix and Perm Buffer with 1ul of Cell-ID Intercalator-Ir. Cells were washed with MaxPar Cell Staining Buffer, resuspended in 1ml of H_2_O/2mM EDTA with 0.5xEQ beads at a concentration of 0.5*10^6^ cells/ml and 35μM-filtered prior to analysis.

### Bulk RNA-sequencing

Cell lines were FACS sorted, washed and resuspended in 1ml of TRIzol™. RNA extraction was performed following manufacturer’s instructions. RNA samples with a RIN ≥ 8 were processed further following Illumina protocols. TruSeq RNA Library Prep Kit (Illumina) was used for samples with high ting cell numbers and 300 ng total RNA was used for library preparation. Libraries were prepared following manufacturer’s instructions with the sole modification of PCR cycle number, which was reduced to 12 instead of the recommended 15 cycles. Libraries were eluted in 30μl of Resuspension Buffer provided with the TruSeq Kit and final concentration was measured by Qubit. 1ng/μl library DNA was analysed on the Bioanalyzer as described above and libraries were diluted based on the corrected Bioanalyzer concentration to 10ng/μl. Libraries were pooled, final pool concentration was re-measured by Qubit and libraries were denatured following Illumina’s protocol. Libraries were diluted to 4nM in 0.2N NaOH and further diluted to 20pM in Tris-HCL, pH7. Libraries were loaded on NextSeq500.

### Bioinformatics

All sequencing data was assessed to detect sequencing failures using *FASTQC* and lower quality reads were filtered or trimmed using *TrimGalore* (https://github.com/FelixKrueger/TrimGalore). Outlier samples containing low sequencing coverage or high duplication rates were discarded.

### ChIP-sequencing analysis

Reads were mapped to the human genome (hg38) using Bowtie (Langmead et al., 2009). Peaks were detected against rabbit IgG control, or in the case of NALM-6 experiments a mock V5 pulldown, using MACS (Zhang et al., 2008). Heatmaps and average signal plots were generated using NGSPlot (Shen, 2014). Read counts were generated for peaks using featureCount*s* from the Rsubread package (Liao et al, 2019). Tracks were visualized using the Integrative Genomics Viewer (IGV, Robinson). Motif enrichment was determined for 501bp windows centred on peak summits using MEME-ChIP (Machanick et al, 2011).

### Processing of RNAseq

Bulk RNAseq samples were mapped to the human reference GRCh38 using tophat2 (https://ccb.jhu.edu/software/tophat/index.shtml). Analyses were performed within the R statistical computing framework, version 4.0.2 using packages from BioConductor version 3.11 (https://Bioconductor.org). Data was combined into a per-gene count matrix using *featureCounts* from the Rsubread package.

### Differential gene expression and principal component analysis

The DEseq2 package (Love et al, 2014) was used for outlier detection, normalization and differential gene expression analyses. Pairwise comparisons were carried out using the Wald test whereas core signatures across multiple cell lines were derived using a likelihood ratio test (LRT) with full model *∼ cell_line + shRNA_target*, reduced model *∼ cell_line*. “Core” RUNX1 target genes (figure 5) are defined as padj<0.1 and direction of change the same in pairwise comparisons (shRNA vs control) for each of the 5 cell lines and LRT padj<0.05. Transformed, normalized counts were obtained by variance stabilizing transformation (VST) for downstream analysis. Principal components were obtained using the *prcomp* function from R base. Plots were generated using the *ggplot2* package (Wickham, 2016) and heatmaps with *pheatmap* (Kolde, 2019).

### Gene sets and Gene Set Enrichment Analysis

Gene signatures of potential biological interest were retrieved from MSigDB version 7.0. In addition, gene signatures derived from label retaining cells were used (Ebinger et al). Gene set enrichment analysis (GSEA) was performed using the R package Cluster Profiler, function GSEA with default arguments. Preranked lists of genes were derived from DESeq2 analysis – genes were ranked according to their Wald-statistic or, in the case of LRT, the rank was calculated as sign(LFC)*-log_10_(pvalue).

### Mass cytometry analysis

Single cell events were gated and exported using Cytobank (https://mrc.cytobank.org/). Clustering (SOM), dimensionality reduction (UMAP) and UMAP plotting were carried out using the R package *CATALYST* (Crowell et al, 2020). Clusters were manually annotated to classify cell types based on cell surface markers, viability based on incorporation of Pt194, cell cycle status based on cell cycle-associated proteins and apoptosis based on expression of cleaved Caspase 3 (Behbehani et al., 2012).

---

**Figure S1.**
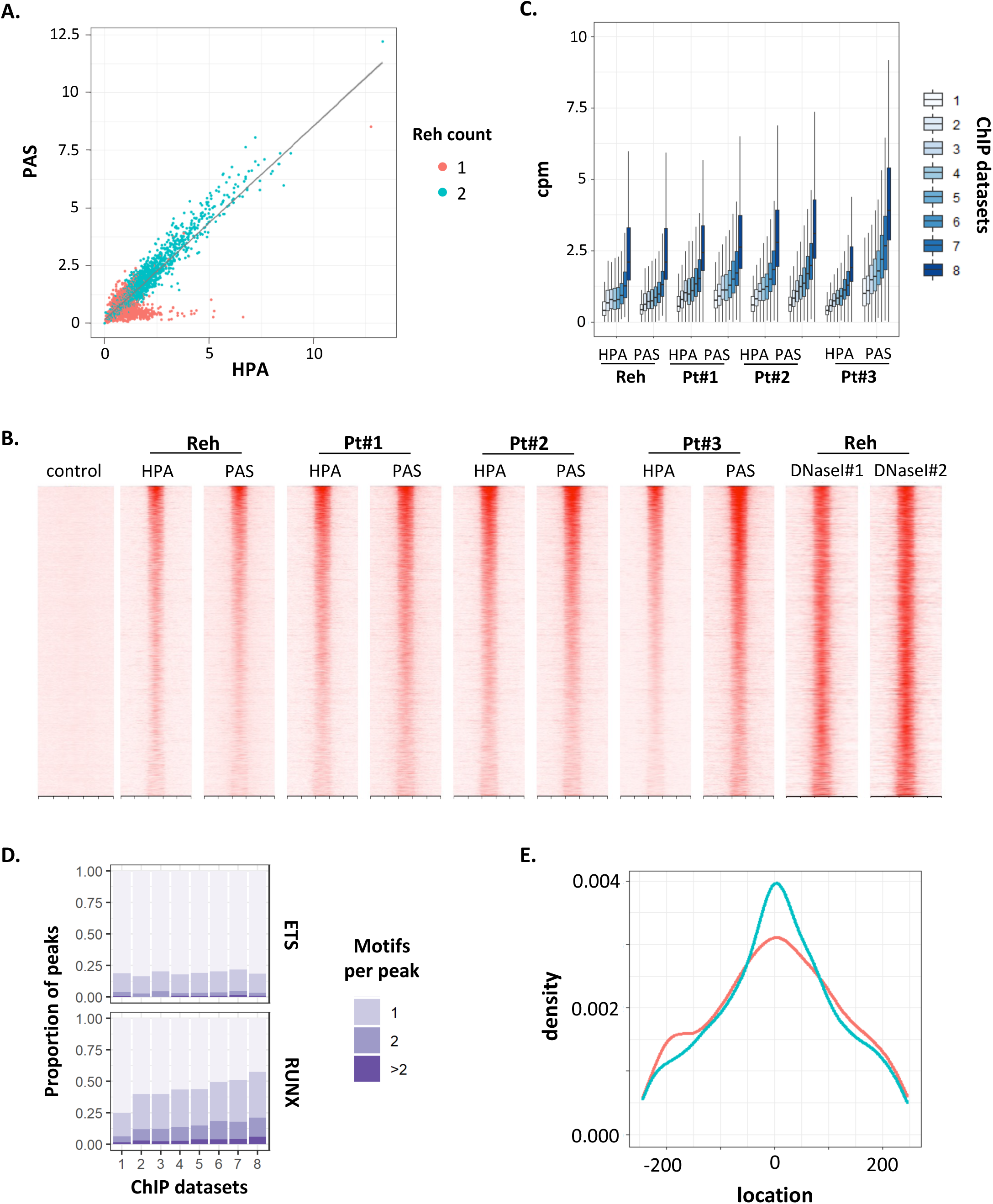
related to Figure 1. (A) Dotplot showing normalised read counts (counts per million) for a 500bp window centred on the summit for all ETV6-RUNX1 peaks identified for two independent ETV6 antibodies (HPA and PAS). Colours represent peaks called in one (1, red dot) or both (2, green dot) ChIP-seq datasets. (B) Heatmap showing ChIP-seq and DNaseI-hypersensitivity read density in a 5kb window centred on peak summits for all ETV6-RUNX1 peaks identified. (C) Boxplot showing distribution of normalised read counts for each of 8 ChIP-seq samples across all ETV6-RUNX1 peaks identified. Colour represents the number of ChIP datasets a peak was identified in. (D) Motif counts in peaks binned according to the number of ETV6-RUNX1 ChIP datasets in which they were identified. (E) Density plot of RUNX and ETS motif distribution across a 500bp window centred on ETV6-RUNX1 ChIP peak summits.

**Figure S2.**
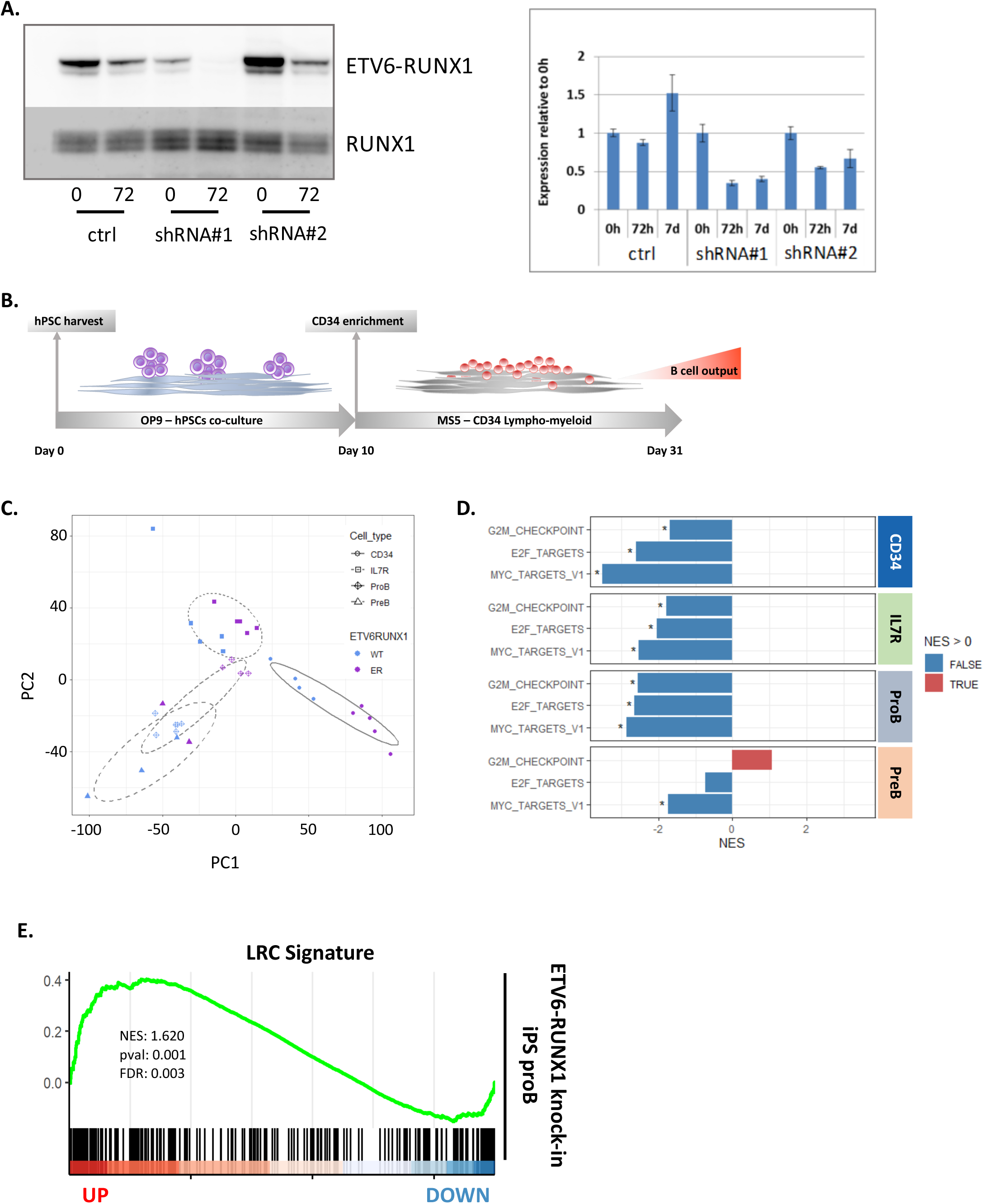
related to Figure 2. (A) Western blot and quantitative PCR showing ETV6-RUNX1 expression in Reh cells for vector control or ETV6-RUNX1 inducible shRNAs 0h, 72h or 7 days (7d) after addition of doxycycline. Error bars indicate standard deviation (n=3). RUNX1 protein levels were unaffected by shRNAs targeting ETV6-RUNX1. (B) Schematic of hPSC B-cell differentiation protocol (C) Principal component analysis of RNA-seq data from wild-type and ETV6-RUNX1 expressing iPSC-derived cell populations (D) Barplot showing normalized enrichment scores (NES) from GSEA for the indicated genesets against pre-ranked gene lists for ETV6-RUNX1 vs wild-type for each of the four populations indicated. * p<0.05 (E) GSEA for label retaining cell (LRC) signature genes against pre-ranked gene list for ETV6-RUNX1 vs wild-type in iPSC-derived proB cells

**Figure S3.**
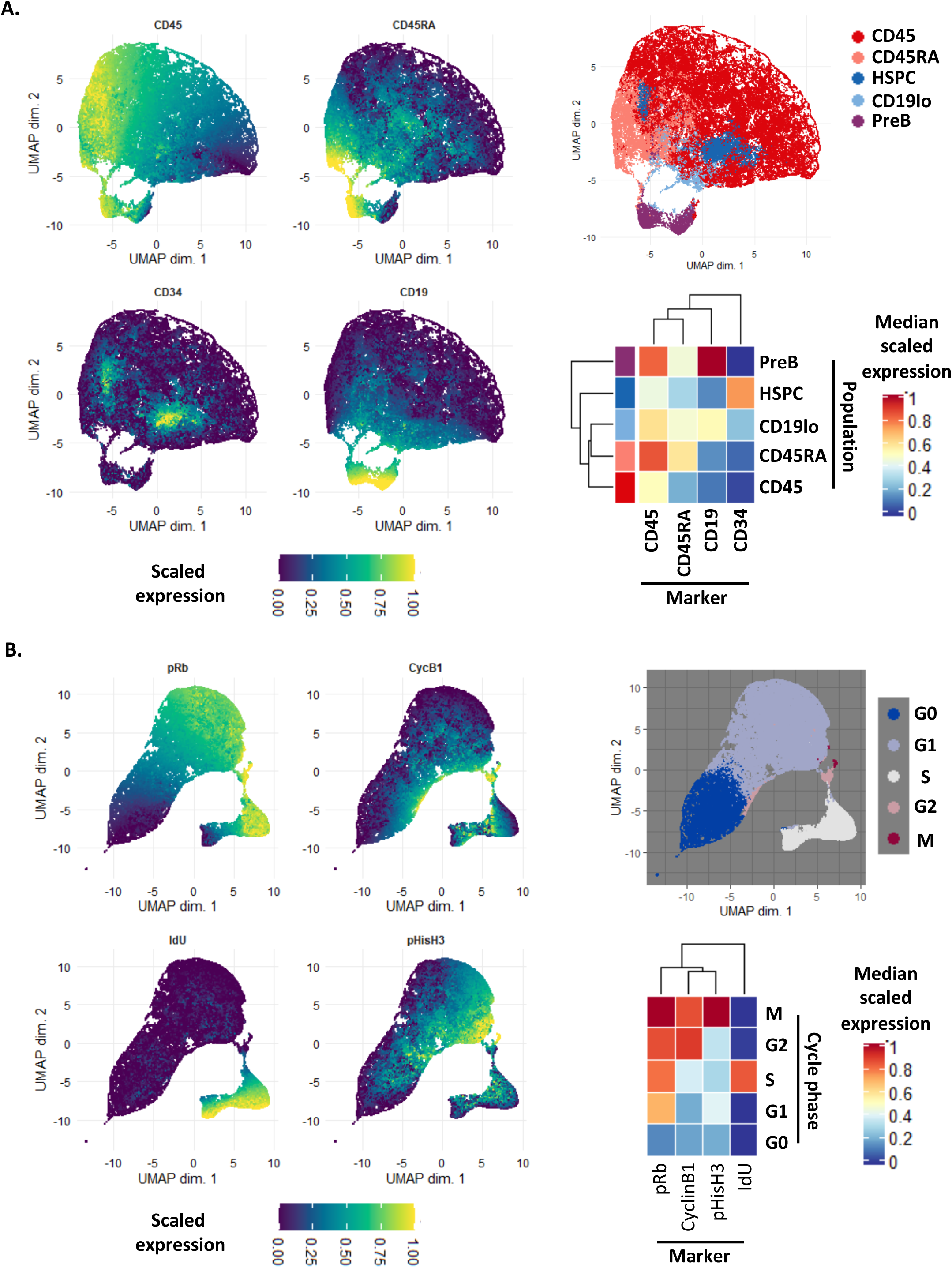
related to Figure 3. (A) UMAP dimensionality reduction for cell surface markers CD45, CD45RA, CD34, and CD19. Colour represents scaled expression of the indicated markers (left) or manually annotated SOM clusters (top right). Heatmap (bottom right) showing median scaled expression of the indicated markers in the annotated clusters. (B) UMAP dimensionality reduction for cell surface markers pRb, IdU, CycB1 and pHisH3. Colour represents scaled expression of the indicated markers (left) or manually annotated SOM clusters (top right). Heatmap (bottom right) showing median scaled expression of the indicated markers in the annotated clusters.

**Figure S4.**
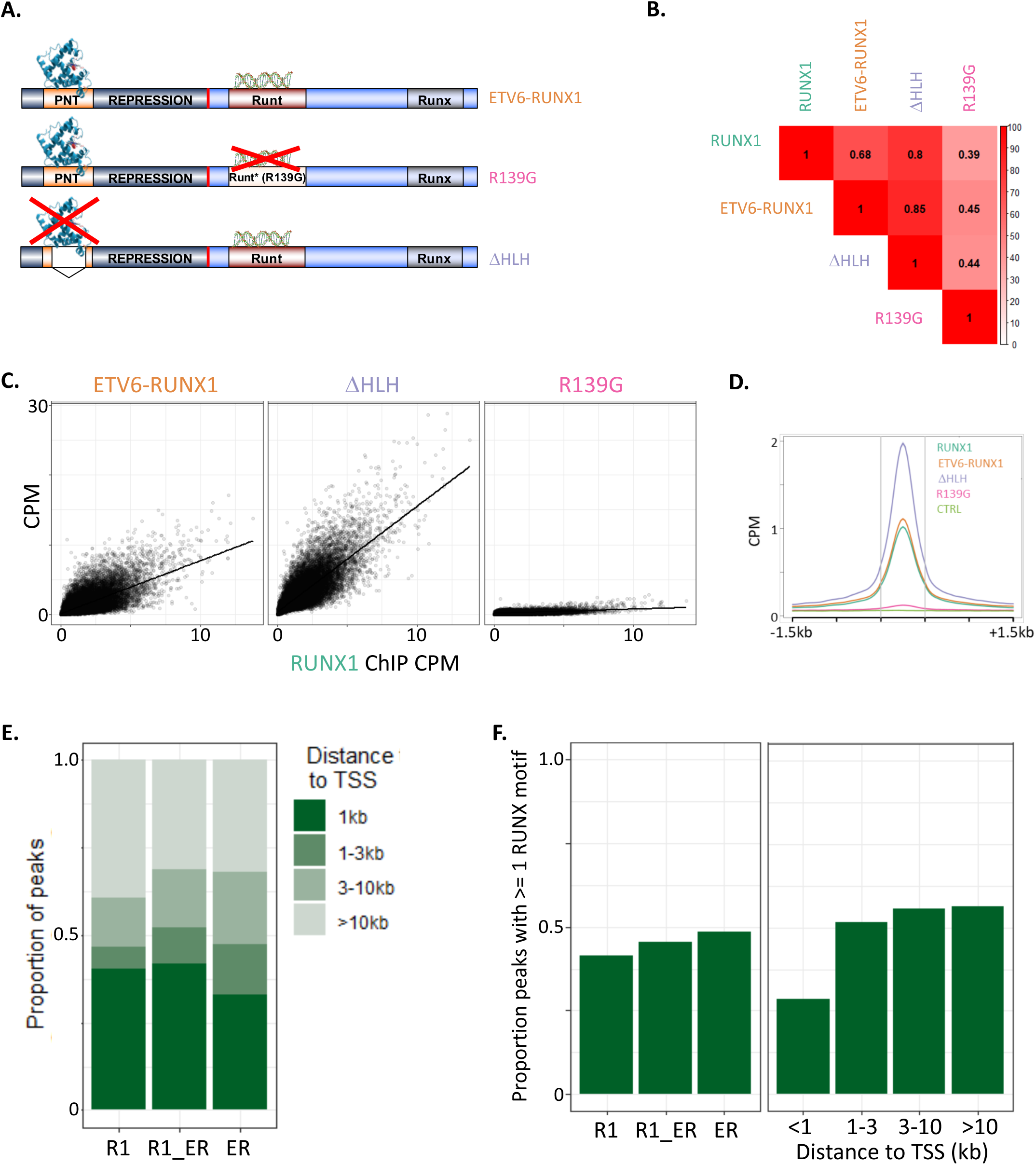
related to Figure 4. (A) Schematic of ETV6-RUNX1 and R139G/ΔHLH mutants. R139G (runt domain point mutation); PNT (pointed domain, required for multimerization). (B) Correlation matrix for ChIP datasets based on normalised read counts (counts per million) for a 500bp window centred on the peak summit across all RUNX1/ETV6-RUNX1 peaks identified. (C) Dotplots of normalised read counts (counts per million (CPM)) for RUNX1 vs each ETV6-RUNX1 dataset. (D) Average profile plot of ChIP-seq signal across all RUNX1/ETV6-RUNX1 peaks identified in a 3kb window centred on peak summits. (E) Proportion of peaks in the indicated bins relative to TSS for each of the peak sets classified in figure 4D. (F) Proportion of peaks with >= 1 RUNX1 motif mapped to a 500bp window centred on the peak summits for peaks binned according to classifications in figure 4D (left panel) or distance from the TSS (right panel).

**Figure S5.**
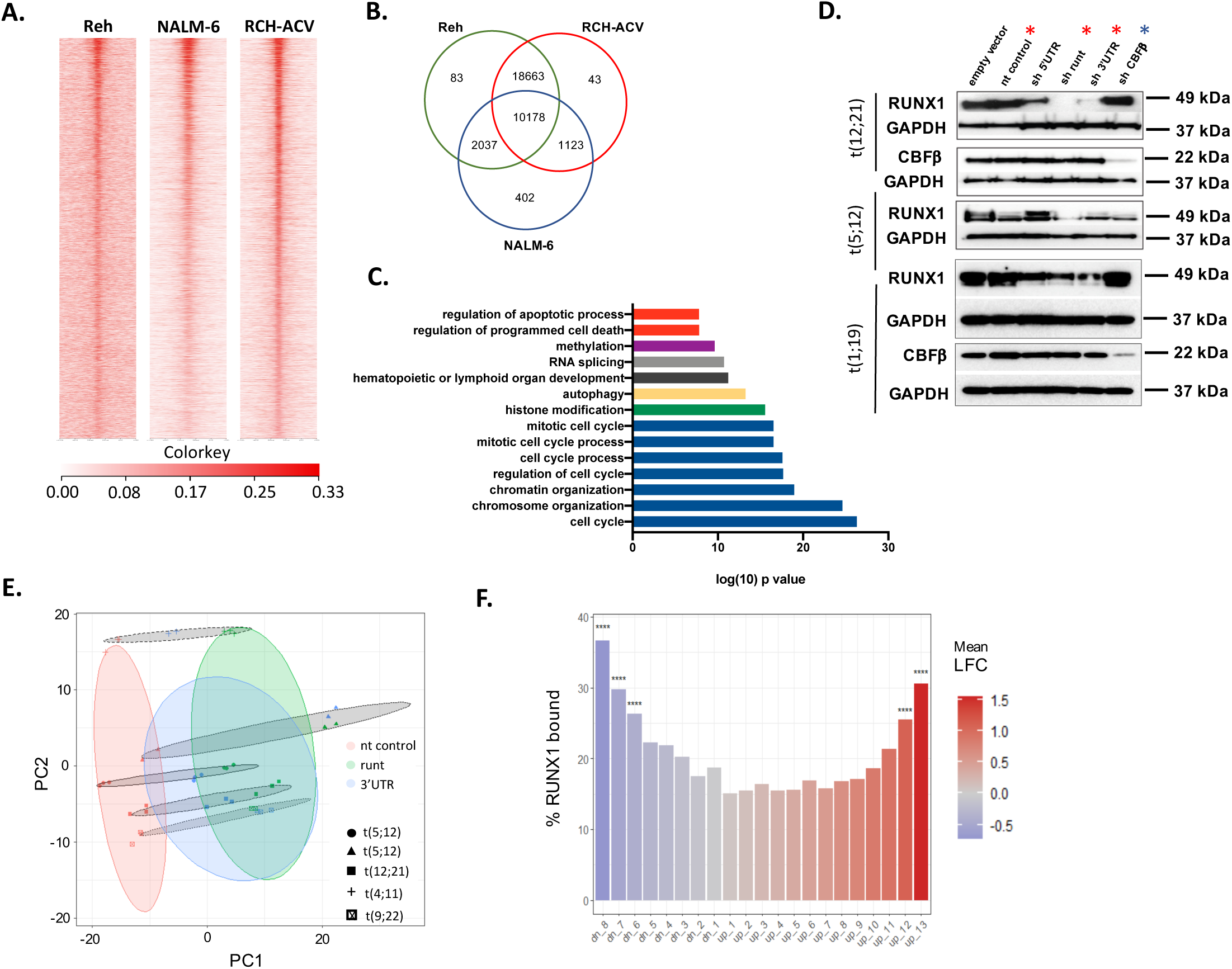
related to Figure 5. (A) Heatmaps showing RUNX1 ChIP-seq signal across all identified RUNX1 binding sites for the indicated cell lines in a 3kb window centred on peak summits. (B) Venn diagram showing overlap of RUNX1 ChIP peaks from Reh, RCH-ACV and Nalm6. (C) Bar plot showing significantly enriched biological processes from GO term analysis (selected biological processes shown). (D) Western blots of Reh, NALM-6 and RCH-ACV cells transduced with control (empty and non-targeting), RUNX1 or CBFβ shRNAs. Protein levels are measured 48h after transduction. Asterisks indicate control and RUNX1 (red) or CBFβ (blue) shRNAs used in subsequent experiments. (E) Dot plot of principal components 1 and 2 showing RUNX1 shRNA and control samples for 5 cell lines. Ellipses show 80% confidence intervals for the indicated groupings. (F) Bar plot showing percentage of genes bound by RUNX1 for genes binned according to response to RUNX1 knockdown. **** p<0.0001, hypergeometric test.

**Figure S6.**
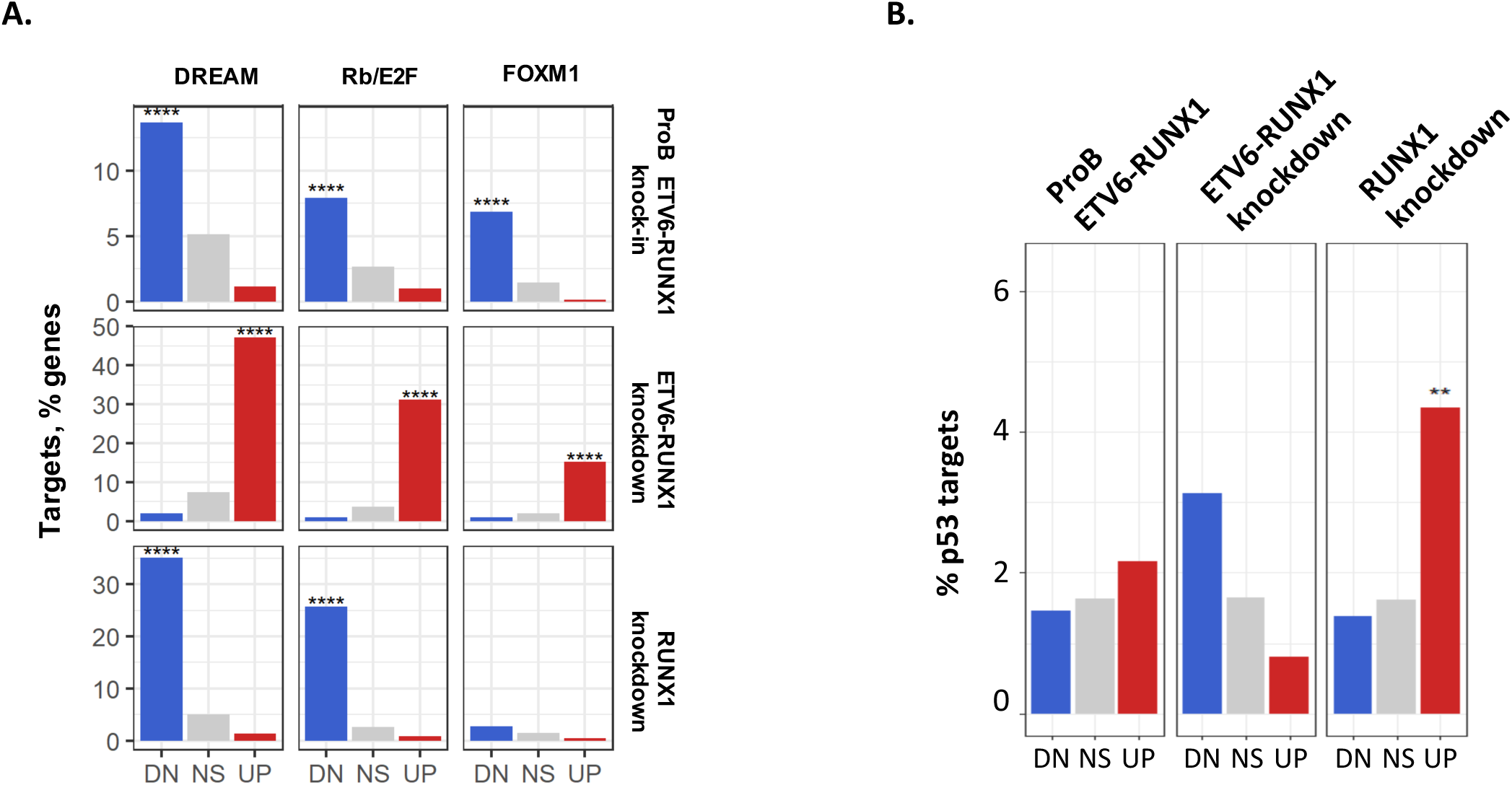
related to Figure 6. (A) Bar plots showing the percentage of genes significantly up-(UP) or down-(DN) regulated in ETV6-RUNX1 knock-in iPSC-derived proB cells, or following ETV6-RUNX1 or RUNX1 knockdown which were defined as DREAM, Rb/E2F, or FOXM1 targets. NS = not significant **** p<0.0001, *** p<0.001, ** p<0.01, * p<0.05, hypergeometric test. (B) Bar plots showing proportion of direct p53 targets (Fischer et al., 2016) in significantly up (UP) or downregulated (DN) genes for ETV6-RUNX1-expressing ProB cells, ETV6-RUNX1 knockdown and RUNX1 knockdown. NS = not significant. **** p<0.0001, ** p<0.01, hypergeometric test

**Figure S7.**
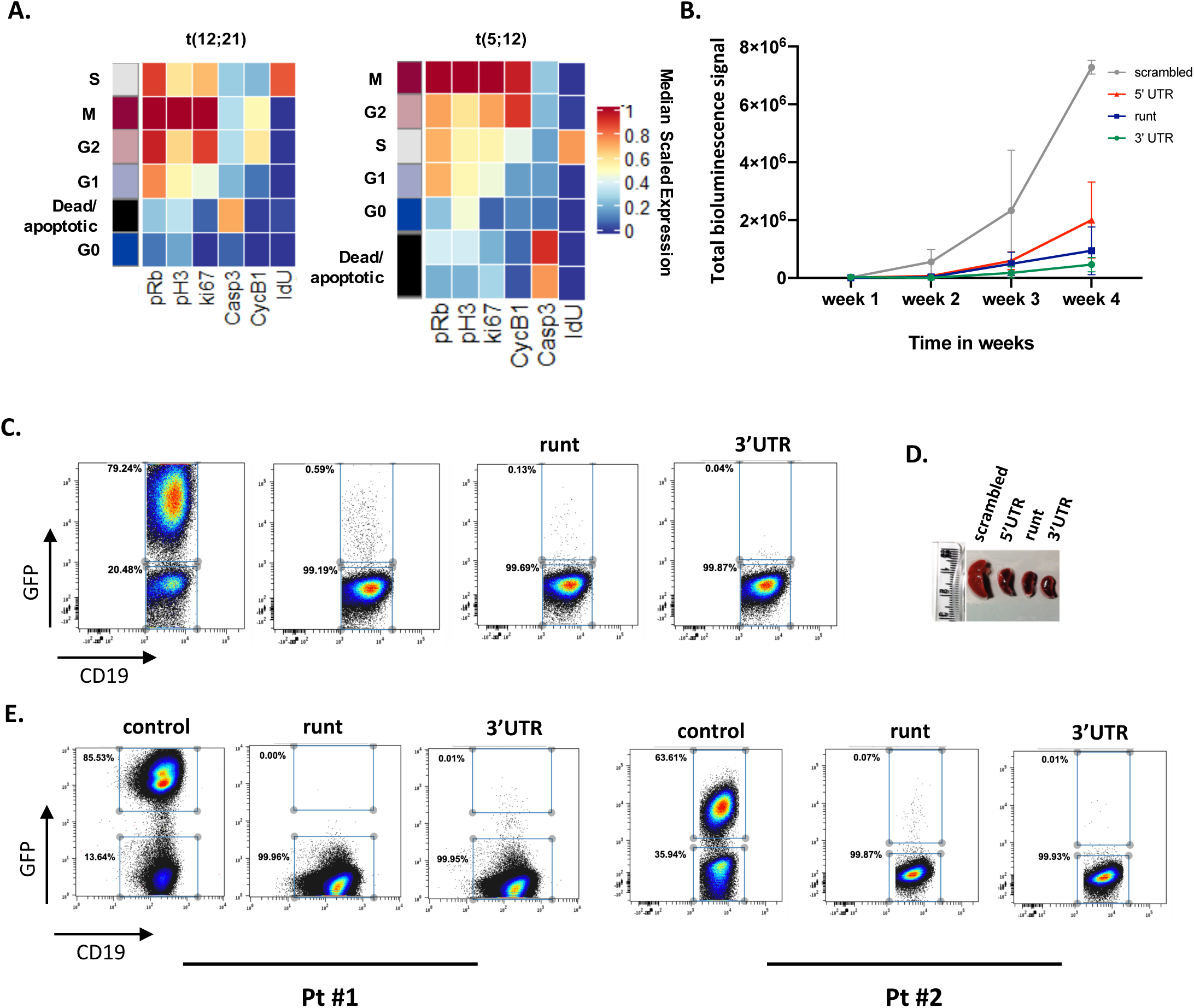
related to Figure 7. (A) Heatmaps showing median scaled expression of the indicated markers in the annotated SOM clusters, corresponding to figure 6B. (B) Summary of bioluminescence imaging of NSG mice engrafted with a 1:1 mix of NALM-6 cells transduced with shRNAs (GFP^+^) and stably expressing a luciferase/RFP reporter were mixed 1:1 with non-transduced (GFP^-^) over four weeks. Relative bioluminescence signal is estimated as the sum of measured signal of ventral and dorsal positions at each time point (n=3). p<0.0001, 2-way Anova test. (C) Representative FACS plot of end point (4 weeks) analysis of NSG mice engrafted with NALM-6 cells as in Figure 7C. (D) Representative image of splenic size obtained from NSG mice engrafted with a 1:1 mix of NALM-6 cells transduced with shRNAs (GFP^+^) and stably expressing a luciferase/RFP reporter were mixed 1:1 with non-transduced (GFP^-^) at week four (end point). (E) Representative FACS plot of endpoint (3 months) analysis of NSG mice competitively engrafted with primary B-ALL cells. Human (CD19^+^) engraftment and the proportion of shRNA-transduced (GFP^+^) cells for two ALL patient samples.

**Figure S8.**
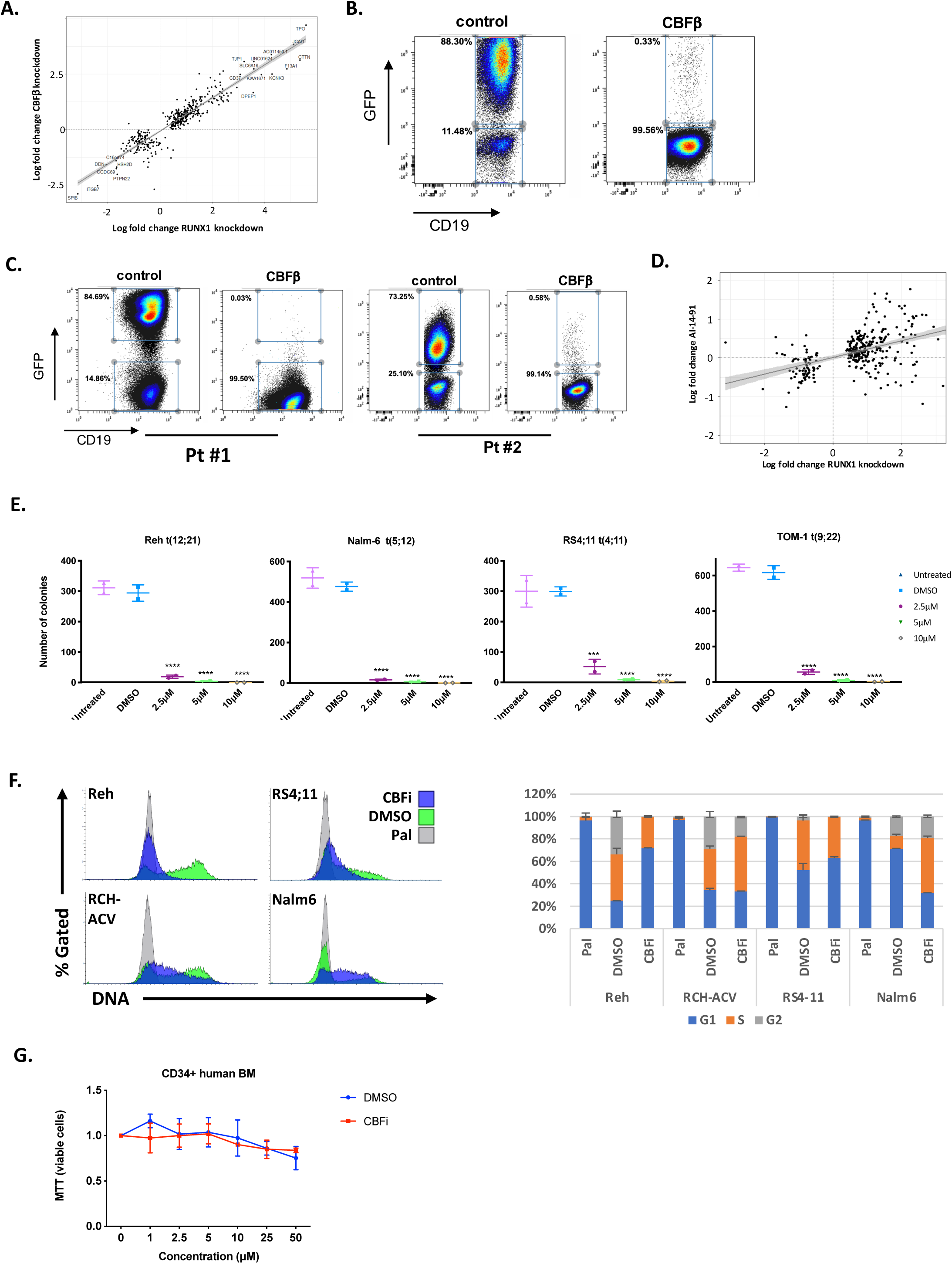
related to Figure 8. (A) Dotplot comparing log fold change following RUNX1 knockdown (x-axis) and CBFβ knockdown (y-axis) in ALL cell lines. “Core” RUNX1 target genes defined in Figure 5 are shown. (B) Representative FACS plot of endpoint (4 weeks) analysis of NSG mice engrafted with NALM-6 cells as in Figure 7C. (C) Representative FACS plot of endpoint (3 months) analysis of NSG mice competitively engrafted with primary B-ALL cells. Human (CD19^+^) engraftment and the proportion of shRNA-transduced (GFP^+^) cells for two ALL patient samples with the indicated genotypes. (D) Dotplot comparing log fold change following RUNX1 knockdown (x-) and CBFi (AI-14-91) treatment (y-axis) in ALL cell lines. “Core” RUNX1 target genes defined in Figure 5 are shown. (E) Colony formation assay of B-ALL cell lines treated with increasing concentrations of CBFi (AI-14-91) (n=2), see Methods for more details. (F) Histograms of DNA content (Hoechst 33342, left panel) showing the impact of AI-14-91 (CBFi) on cell cycle re-entry following palbociclib (Pal) wash-out. Following wash-out cells were incubated for a further 18h in the presence of Pal, CBFi or DMSO. Quantification (right panel) of cell cycle distribution based on DNA content (G1=2N, 2N<S<4N, G2=4N). Error bars indicate standard deviation (n=3). Note that Nalm6 transit through the cell cycle more rapidly, hence a high proportion of control (DMSO) cells have re-entered G1. (G) Plot showing sensitivity of CD34^+^ human bone marrow (hBM) cells to AI-14-91 (CBFi). Cells were cultured in the indicated concentrations of CBFi or vehicle control (DMSO) for 48h and MTT assay for cell viability performed. Values were normalised to untreated (0μM, DMSO) cells. Error bars indicate standard deviation (n=3).

## Notes

### Competing Interest Statement

The authors have declared no competing interest.

